# Glial alterations in the glutamatergic and GABAergic signaling pathways in a mouse model of Lafora disease, a severe form of progressive myoclonus epilepsy

**DOI:** 10.1101/2024.09.13.612874

**Authors:** Rosa Viana, Teresa Rubio, Ángela Campos-Rodríguez, Pascual Sanz

## Abstract

Lafora disease (LD; OMIM#254780) is a rare form of progressive myoclonus epilepsy characterized by the accumulation of insoluble deposits of aberrant glycogen (polyglucosans), named Lafora bodies (LBs), in the brain but also in peripheral tissues. It is assumed that the accumulation of LBs is related to the appearance of the characteristic pathological features of the disease. In mouse models of LD, we and others have reported an increase in the levels of reactive astrocytes and activated microglia, which triggers the expression of the different pro-inflammatory mediators. Recently, we have demonstrated that the TNF and IL-6 inflammatory signaling pathways are the main mediators of the neuroinflammatory phenotype associated with the disease. In this work, we present evidence that the activation of these pathways produces a dysregulation in the levels of different subunits of the excitatory ionotropic glutamatergic receptors (phopho-GluN2B, phospho-GluA2, GluK2) and also an increase in the levels of the GABA transporter GAT1 in the hippocampus of the *Epm2b-/-* mice. In addition, we present evidence of the presence of activated forms of the Src and Lyn protein kinases in this area. These effects may increase the excitatory glutamatergic signaling and decrease the inhibitory GABAergic tone, leading to hyper-excitability. More importantly, the enhanced production of these subunits occurs in non-neuronal cells such as activated microglia and reactive astrocytes, pointing out a key role of glia in the pathophysiology of LD.

## INTRODUCTION

Lafora disease (LD; OMIM#254780) is a rare form of progressive myoclonus epilepsy characterized by the accumulation of insoluble deposits of aberrant glycogen (polyglucosans), named Lafora bodies (LBs), in the brain but also in peripheral tissues [1]. LD clinical manifestations appear during childhood or early adolescence. Clinically, LD shares some symptoms with other progressive myoclonus epilepsies (EPMs), such as generalized tonic-clonic seizures, myoclonus, absences, and visual hallucinations. However, LD is the most severe form of EPM since patients present a rapid deterioration and dementia with amplification of seizures, leading to death after around a decade from the onset of the first symptoms [2], [3]. LD is caused by mutations in two genes, namely *EPM2A*, encoding the glucan phosphatase laforin, and *EPM2B*, encoding the E3-ubiquitin ligase malin [4], [5]. Both proteins form a functional complex where laforin recognizes possible substrates that are then ubiquitinated by malin [6]. The laforin/malin complex regulates negatively glycogen synthesis by modulating the activity of glycogen synthase and also by regulating the uptake of glucose [7], [8]. This could explain why glycogen synthesis is enhanced in LD patients, leading to the accumulation of LBs. In addition, the laforin/malin complex is also involved in the regulation of other physiological pathways such as the maintenance of proteostasis and the response to oxidative stress [2], [9].

Mouse models have been used to study the molecular basis of LD pathophysiology: *Epm2a-/-* mice, which lack exon 4 from the *Epm2a* gene [10]; and *Epm2b-/-* mice, which lack the single exon present in the *Epm2b* gene [11]. Both mouse models present similar pathophysiological phenotypes, which resemble the ones present in LD patients: e.g., they show an accumulation of LBs in the brain and peripheral tissues [11]; they are more sensitive to the effects of pro-convulsant drugs such as kainate or pentylenetetrazole (PTZ) [12]; and they exhibit a decreased ability to glutamate uptake, which could lead to excitotoxicity [13]. In addition, they show signs of neuroinflammation, with an increase in the levels of reactive astrocytes and activated microglia, which trigger the expression of the different pro-inflammatory mediators [14]. Recently, we have determined that neuroinflammation is mediated by the activation of two main inflammatory signaling pathways, namely TNF and IL-6, and that there is an infiltration of peripheral cytotoxic T-lymphocytes in the brain parenchyma of *Epm2b-/-* mice [15].

It is important to bear in mind that the activation of the TNF signaling pathway contributes to neuronal hyper-excitability [16] because, on the one hand, it increases the levels of glutamate in the brain by reducing the activity of glutamate transporters that are involved in glutamate uptake, and also by enhancing the release of this neurotransmitter from the astrocytes by improving the function of the xCT cystine-glutamate antiporter [17], [18]. On the other hand, TNF signaling increases the functionality of the post-synaptic excitatory glutamatergic NMDA and AMPA receptors and also induces the endocytosis of post-synaptic inhibitory GABA receptors [16]. In summary, activation of the TNF signaling pathway leads to hyper-excitability by an increase in the excitatory and a decrease in the inhibitory neurotransmission [19], [20]. In this work, we studied whether in an animal model of LD (*Epm2b-/-* mice) there were alterations in the glutamatergic and GABAergic signaling pathways which may account for the hyper-excitability observed in these animals [12]. With this aim, we analyzed the main ionotropic (NMDA, AMPA, and Kainate) and metabotropic (mGluRs) excitatory glutamatergic receptors, and also the inhibitory GABA receptors and the GABA transporters (GATs).

The NMDA (N-methyl-D-aspartate) glutamate receptors are tetramers composed of 2 GluN1 subunits and combinations of 2 GluN2 or GluN3 subunits. The GluN1 and GluN3 subunits bind the co-agonists glycine and D-Serine, whereas the GluN2 subunit is in charge of binding glutamate. They are mainly expressed and located at the post-synaptic neuronal terminals [21], and upon binding to glutamate, NMDA receptors allow the entry of Na+ and Ca++. In a physiological setting, Ca++ signaling through NMDA receptors maintains a defined transcriptional program, and regulates synaptic strength, memory consolidation, and neuronal survival. However, when overactivated, NMDA receptors allow the entry of excess of Ca++, which is detrimental and causes excitotoxicity leading to neuronal demise [22]. The activity of these NMDA receptors is modulated by post-translational modifications (e.g., phosphorylation, ubiquitination). For example, under conditions of neuroinflammation, the activation of the TNF, IL-6, and IL-1b signaling cascades leads to an increase in the activity of the NMDA receptors, since in the short run, there is an activation of the Src family of protein kinases which phosphorylate NMDA subunits [23], [24], improving in this way the recycling of the subunits to the plasma membrane; this increases the entry of Ca++ leading to a reduction of the excitatory threshold [16], [25]. In the long run, these signaling events increase the expression of genes related to epilepsy [16], [26], [27]. Astrocytes and microglia also express NMDA receptors; they participate in glutamate signaling and increase the entry of Ca++ in these cells [28], [29].

The AMPA (α-amino-3-hydroxy-5-methyl-4-isoxazolepropionic acid) glutamate receptors are tetramers composed of combinations of GluA1-GluA4 subunits. The GluA1 and GluA2 are the most important subunits and form a typical 2GluA1-2GluA2 receptor. They are mainly expressed and located at the post-synaptic neuronal terminals [21]. Upon binding of glutamate, AMPA receptors mediate the entry of Na+, which produces the depolarization of the post-synaptic membrane, an event that is necessary to release Mg++ from its binding to the NMDA receptor, allowing in this way its activation [30], [31]. The activity of the AMPA receptors is regulated by post-translational modifications (e.g., phosphorylation, ubiquitination), which controls their recycling from the plasma membrane to endosomal compartments and also mediates their degradation. Some subunits of the AMPA receptors (e.g., GluA2) are also expressed in astrocytes and microglia [21], [28], [32].

The Kainate glutamate receptors are tetramers composed of GluK1-GluK5 subunits. They are mainly expressed and located at the post-synaptic neuronal terminals [21]. GluK2 and GluK5 form a Kainate receptor that participates in the generation of seizures in patients suffering from temporal lobe epilepsy (TLE) [33]. It has also been reported a direct participation of receptors containing the GluK2 subunit in epilepsy [30]. GluK2 is expressed in hippocampal pyramidal cells (CA1 y CA3), hippocampal dentate gyrus (DG), cerebellar granule cells, and cortical pyramidal cells, whereas GluK5 is expressed abundantly in all brain areas [34]. In addition, some subunits of the Kainate receptors (e.g., GluK2, GluK5) are also expressed in astrocytes [21], [28].

Finally, monomeric metabotropic glutamate receptors (mGluRs) also participate in glutamate signaling at the post-synaptic neuronal site. Among the different mGluRs, mGluR3 and mGluR5 have a major role. mGluR5 receptor interacts with post-synaptic NMDA receptors and improves their activity. In addition, mGluR3 and mGluR5 are expressed in astrocytes [21] and regulate Ca++ entry in these cells, initiating the production of arachidonic acid followed by the formation of vasoactive substances [19], [29], [30].

On the other hand, the inhibitory GABAergic signaling pathway is dependent on the presence of ionotropic GABAA and metabotropic GABAB receptors, and also on the presence of GABA transporters (GATs). GABAA receptors are heteropentamers composed of 2 alpha, 2 beta, and 1 gamma subunit. At the hippocampus, the most abundant GABAA receptor is formed by alpha1 and gamma2 subunits [23]. In addition to inhibitory neurons, astrocytes also express GABAA and GABAB receptors, as well as GABA transporters (mainly GAT1 and GAT3). Therefore, astrocytes can respond and control the extracellular levels of GABA [35].

In this work, we present evidence of a dysregulation in the levels of different subunits of the excitatory ionotropic glutamatergic receptors and also an increase in the levels of the GABA transporter GAT1 in the hippocampus of the *Epm2b-/-* mice. In addition, we present evidence of the presence of activated forms of the Src and Lyn protein kinases in this area. These effects may increase the excitatory glutamatergic signaling and decrease the inhibitory GABAergic tone, leading to hyper-excitability. More importantly, the enhanced production of these subunits occurs in non-neuronal cells such as activated microglia and reactive astrocytes, pointing out a key role of glia in the pathophysiology of LD.

## MATERIAL AND METHODS

### Ethics statement, animal care, mice and husbandry

This study was carried out in strict accordance with the recommendations in the Guide for the Care and Use of Laboratory Animals of the Consejo Superior de Investigaciones Científicas (CSIC, Spain) and approved by the Consellería de Agricultura, Medio Ambiente, Cambio Climatico y Desarrollo Rural from The Generalitat Valenciana. All procedures were approved by the animal committee of the Instituto de Biomedicina de Valencia, CSIC (authorization number IBV-51). All efforts were made to minimize animal suffering. Although it has been described that in LD there are no differences in terms of pathophysiology between males and females, in this study we used both sexes of homozygous *Epm2b-/-* mice in a pure C57BL/6JRccHsd background and the corresponding wild type (control) of 16 months of age. Mice were maintained in the IBV-CSIC facility on a 12/12 light/dark cycle under constant temperature (23°C) with food and water provided *ad libitum*. For the immunofluorescence experiments, at least four individuals (including males and females) from each genotype were used in this study. They were perfused as described below. An additional three individuals from each group were used for the Western blot experiments. When planning the experiments, the principles outlined in the ARRIVE guidelines and the Basel declaration including the 3R concept have been considered.

### Preparing brain tissue by conventional cardiac perfusion

The animals were deeply anesthetized with sodium pentobarbital 80 mg/Kg by intraperitoneal injection and perfused via the heart, using a perfusion pump, at a speed of 8 ml/min with 4% paraformaldehyde (PFA), in PBS (40-48 ml per animal). At the beginning, blood removal from the circulatory system was performed with 0.9% saline solution (24 ml per animal) before the fixative entered. After perfusion, brains were removed and incubated in 15 ml of 4% PFA at 4°C with shaking up for 2h. Then, they were washed three times with PBS at room temperature for 10 min. Next, brains were incubated for 30 min with 50% ethanol, and then they were incubated overnight in 70% ethanol. The next day, the two hemispheres were separated, dehydrated, cleared, and embedded in paraffin.

### Immunofluorescence analyses

At least four individuals (including at least 2 males and 2 females) from each genotype of 16-month-old mice were perfused as described above. The samples embedded in paraffin were sagittally sectioned at 4 μm using a microtome HistoCore Biocut (Leica, Madrid, Spain). Sections corresponding to similar brain coordinates were deparaffined and rehydrated, and then, microwave antigen retrieval was performed for 10 min in 10 mM citrate buffer pH 6.0, or 10 mM Tris 1 mM EDTA pH 9.0, following the antibody manufacturer’s instructions in each case. Sections were immersed in blocking buffer (1% BSA, 10% FBS, 0.2% Triton X-100, in PBS) and incubated overnight at 4°C with the corresponding primary antibodies listed in Supplementary Table S1 diluted in blocking buffer. Some primary antibodies required a pre-incubation with a 400 mM glucose solution at 4°C overnight to avoid cross-reaction with polyglucosan bodies [36]. After three washes of 10 min in PBS, sections were incubated for one hour at room temperature with the appropriate secondary antibody (Supplementary Table S1) diluted at 1/500 in blocking buffer without Triton X-100, washed once with PBS, incubated with DAPI (Sigma, Madrid, Spain), washed twice with PBS and mounted in AquaPolymount (Polysciences Inc., USA). Images were acquired in a Confocal Spectral Zeiss LSM 980 microscope (Zeiss group, Germany). Pictures of each area of the whole hippocampus were taken with a 40x objective. All images were acquired at one unit Airy pinhole, with a resolution of 512 × 512 pixels and z stacks of 0.8 μm. Image stitching and maximum projection of all the images were performed using the ZEN Blue (Zeiss) software. At least four independent samples were analyzed for each staining and condition (control and *Epm2b-/-* mice). For each staining, the percentage of the corresponding laser intensity and detector gain settings were maintained between samples. For image analysis, the background obtained using only a secondary antibody was subtracted using the image-processing package Fiji-ImageJ, and the intensity of the signal was normalized by the size of the studied area, as in [15]. These areas within the hippocampus were: CA1 and CA3, pyramidal layers of the Cornu ammonis 1 and 3, respectively; DG, the granular layer of the dentate gyrus; and SR, the stratum radiatum area between the CA1 and DG regions, containing the radiatum, the lacunosum-moleculare, and the molecular layers (Supplementary Figure S1A). As we did not detect differences between male and female samples, we collected all their signals for statistical analyses. Amplified photos of particular areas of the hippocampus were obtained using the 63x objective, with a resolution of 1024 × 1024 pixels and z stacks of 0.5 μm. In colocalization experiments, the selected area was the same for all channels in the same image.

### Western blot and immunodetection

Mice used for western blot analyses (three animals from each group, including males and females) were sacrificed by cervical dislocation, brains were rinsed twice with cold phosphate buffer saline (PBS) (0.1 M, pH 7.4), and hippocampi rapidly separated and frozen immediately in liquid nitrogen. Mouse brain hippocampi were lysed in RIPA buffer [50 mM Tris-HCl, pH 8; 150 mM NaCl; 0.5% sodium deoxycholate; 0.1% SDS; 1% Nonidet P40; 1mM PMSF; and complete protease inhibitor cocktail (Roche, Barcelona, Spain)] for 30 min at 4°C with occasional vortexing. The lysates were passed ten times through a 25 gauge needle in a 1 ml syringe and centrifuged at 13,000 x g for 15 min at 4°C. Supernatants were collected and a total of 35 μg protein was subjected to SDS-PAGE and transferred onto a PVDF membrane. Membranes were blocked in 5% (w/v) nonfat milk in Tris-buffered saline (TBS-T: 50 mM Tris-HCl, 150 mM NaCl, pH 7.4; with 0.1% Tween-20) for 1 h at room temperature and incubated overnight at 4°C with the corresponding primary antibodies listed in Supplementary Table S1. Mouse anti-GAPDH (Santa Cruz Biotechnologies, sc-32233) was used as a housekeeping antibody. Then, membranes were probed with suitable secondary antibodies for 1 h at room temperature. Signals were obtained by chemiluminescence using ECL Prime Western Blotting Detection Reagents (Cytiva-Amersham, RPN2232), and the image reader Fuji-LAS-4000 (GE Healthcare, Barcelona, Spain). The results were quantified using the software Image Studio Lite version 5.2 (LI-COR Biosciences, Germany). Related quantifications of the detected proteins to the levels of GAPDH are shown in Supplementary Table S2.

### Statistical analyses

Experiments were performed in three mice from each genotype for the Western blot analyses and at least four mice (males and females) for immunofluorescence analyses. The data was previously analyzed for normal (Gaussian) distribution by D’Agostino-Pearson Omnibus normality as well as Shapiro-Wilk normality tests, using GraphPad Prism version 6.0 statistical software (La Jolla, CA, USA). The alpha value was less than 0.05 for both tests, meaning that the data did not follow a normal Gaussian distribution. Consequently, an unpaired and non-parametric Mann-Whitney test was performed, according to GraphPad software. Values are expressed as median with a range ± standard deviation. P-values have been considered as *p<0.05; n: 3 independent samples in the Western blot analyses and at least four or more in the immunofluorescence analyses.

## RESULTS

In this work, we have analyzed the levels of different subunits of the glutamatergic and GABAergic signaling pathways in the brain of LD animals (*Epm2b-/-*) of 16 months of age and compared them to control animals of the same age, including males and females of both genotypes.

### 1. Glutamatergic signaling

To study the possible alterations in glutamatergic signaling, we analyzed the levels of several subunits of the different glutamatergic receptors (NMDA, AMPA, Kainate, and mGluR), in the brain of LD and control animals by immunofluorescence using specific antibodies (Supplementary Table S1). We focused our attention on the hippocampus since it has been reported that this is the region that is mainly related to hyper-excitability [37]. We analyzed 4 areas in the hippocampus: Cornu ammonis 1 (CA1), Cornu ammonis 3 (CA3), Dentate gyrus (DG), and Stratum radiatum (SR), the region between CA1 and DG regions, containing the radiatum, the lacunosum-moleculare, and the molecular layers (see Supplementary Fig. S1A).

#### 1.1. NMDA receptor subunits

We analyzed the levels of the most important post-synaptic subunits of the NMDA receptor, namely GluN1, GluN2A, and GluN2B [19], [38]. In the case of GluN1 and GluN2A subunits, when we analyzed the whole hippocampus, we detected a weak signal above the basal intensity obtained when using only the secondary antibody, but no differences in intensity between control and *Epm2b-/-* samples were observed in the 4 areas indicated above (Supplementary Fig. S1B). Then, we co-labelled the selected NMDA subunits with vGlut1 or synaptophysin (pre-synaptic neuronal markers) and analyzed a higher magnification of these areas. In this way, we wanted to assess whether there were changes in the subcellular distribution of the NMDA subunits, as it has already been reported to occur under conditions of status epilepticus [39]. In the control samples, the anti-GluN1 and anti-GluN2A antibodies labeled the pyramidal neurons of the CA1 region as well as the granular neurons of the DG region, with an additional punctuated nuclear staining in the case of GluN2A (Fig. 1A and 1B, in green). The same was true for the CA3 region (not shown). No changes in the subcellular distribution of any of these subunits were detected in the *Epm2b-/-* samples in these areas. Nevertheless, in the case of GluN2A staining, we observed the appearance of more dysmorphic neurons in the CA1 and DG regions of LD samples (Fig. 1B; white arrows), which could be a sign of altered neuronal functionality. The appearance of dysmorphic neurons might confirm the reported alterations in neuronal connectivity present in animal models of LD [40]. When we analyzed the SR region, no major differences between LD and control samples were observed either (Fig. 1A and 1B). Western blot analysis of whole hippocampi did not show changes in the levels of the GluN2A subunit; unfortunately, no signal was detected with the GluN1 antibody (Supplementary Fig. S9A).

**Figure 1:**
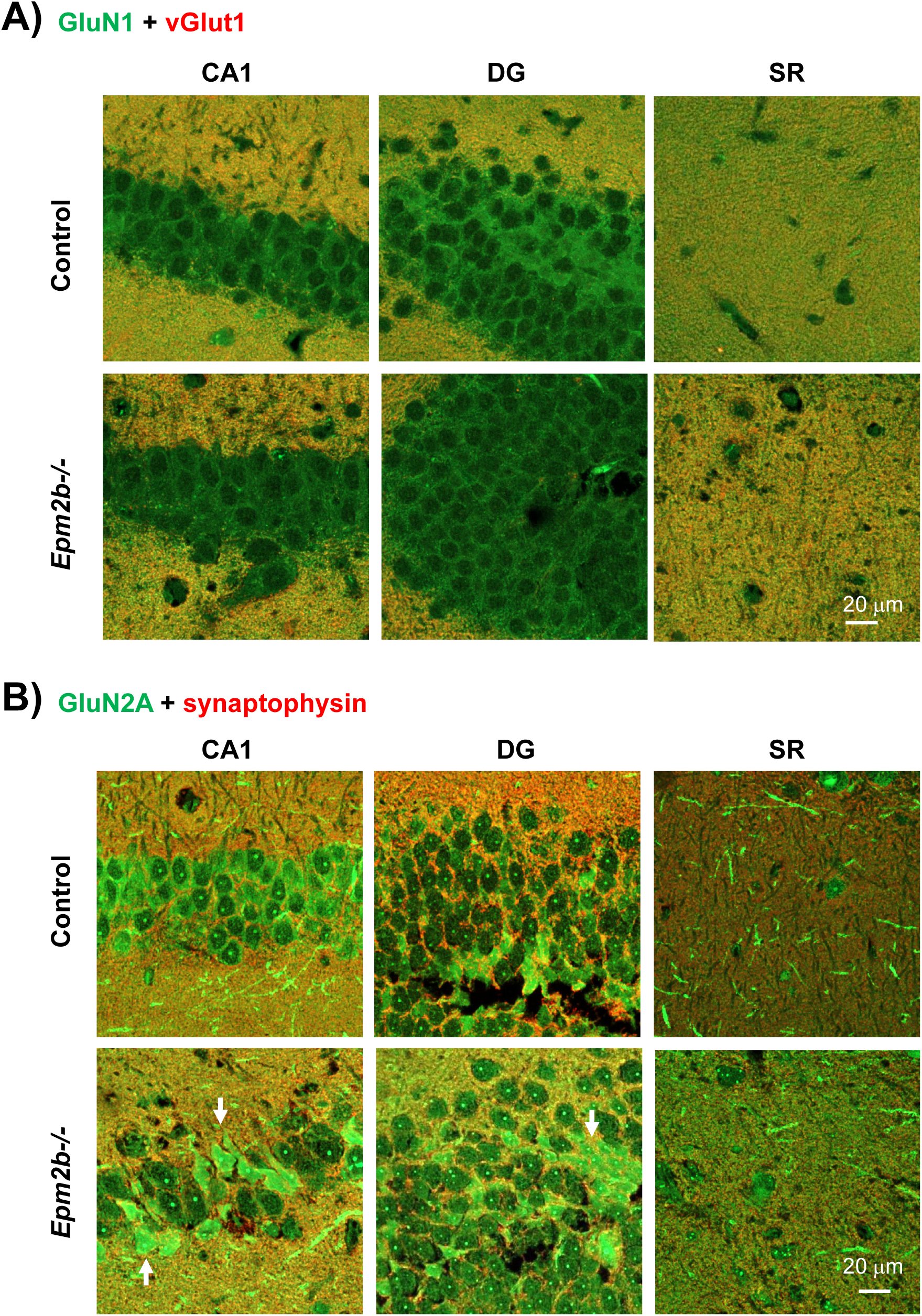
Immunofluorescence analyses of the GluN1 and GluN2A subunits of the NMDA receptor. A magnification of the Cornu ammonis (CA1), Dentate gyrus (DG), and Stratum radiatum (SR; containing the radiatum, the lacunosum-moleculare, and the molecular layers) regions of representative confocal images of the hippocampus derived from control and *Epm2b-/-* mice of 16 months of age, co-labelled by immunofluorescence with anti-GluN1 (A) or anti-GluN2A (B) (in green) and the pre-synaptic markers anti-vGlut1 (in A) or anti-synaptophysin (in B) (in red) antibodies, is shown. At least four independent samples from each genotype (males and females) were analyzed in the same way. The scale corresponds to 20 micrometers. In panel B), white arrows indicate dysmorphic neurons.

We also analyzed the levels of the GluN2B subunit at the hippocampal level but we could not reach any conclusion since the antibody we used cross-reacted with the Lafora bodies present in the LD samples, even when pre-incubated with 400 mM glucose to avoid cross-contamination [36] (Supplementary Fig. S1B). Western blot analysis did not show differences between control and LD samples (Supplementary Fig. S9A). Since it has been described that the GluN2B subunit is phosphorylated by members of the Src family of protein kinases as a result of TNF activation, we checked the presence of phosphorylated forms of the GluN2B subunit (pGluN2B-Y1336), alone (Fig. 2A), or in combination with synaptophysin (Fig. 2B). We observed an increase of the intensity of the pGluN2B signal but only in the SR region of the hippocampus of *Epm2b-/-* samples in comparison to controls (13.37 ± 2.58 a.u. in control vs 17.41 ± 2.19 a.u. in *Epm2b-/-* samples; p-value: 0.0381) (Fig. 2A; Supplementary Table S2); we did not observe major changes in the intensity of the neuronal staining of the CA1, CA3, and DG areas between control and *Epm2b-/-* samples (not shown). The higher signal of pGluN2B in the SR area of *Epm2b-/-* samples corresponded to cellular structures not present in the control samples (Fig. 2B). To elucidate which cells expressed the pGluN2B signal, we performed a co-staining with GFAP (glial fibrillary acidic protein; an astrocytic marker) and Iba1 (ionized calcium-binding adaptor molecule 1; a microglial marker). In Fig. 3A, we show an increase of the GFAP and Iba1 staining in the LD samples as already reported [15]. The signal of pGluN2B in the SR area corresponded to microglia and probably infiltrated lymphocytes present in the brain parenchyma of LD animals [15], but not to astrocytes. This conclusion is supported by a magnification of the SR area of LD samples presented in Fig. 3B. Clearly, cells expressing the Iba1 marker (microglia) also expressed the pGluN2B marker. Western blot analysis did not shoew differences between control and LD samples (Supplementary Fig. S9A); perhaps, as the samples prepared for Western blot contained the whole hippocampus, the neuronal signal could have masked the signal of the SR region. In any case, our results indicate that in LD samples, microglial cells express the pGluN2B subunit of the NMDA receptor.

**Figure 2:**
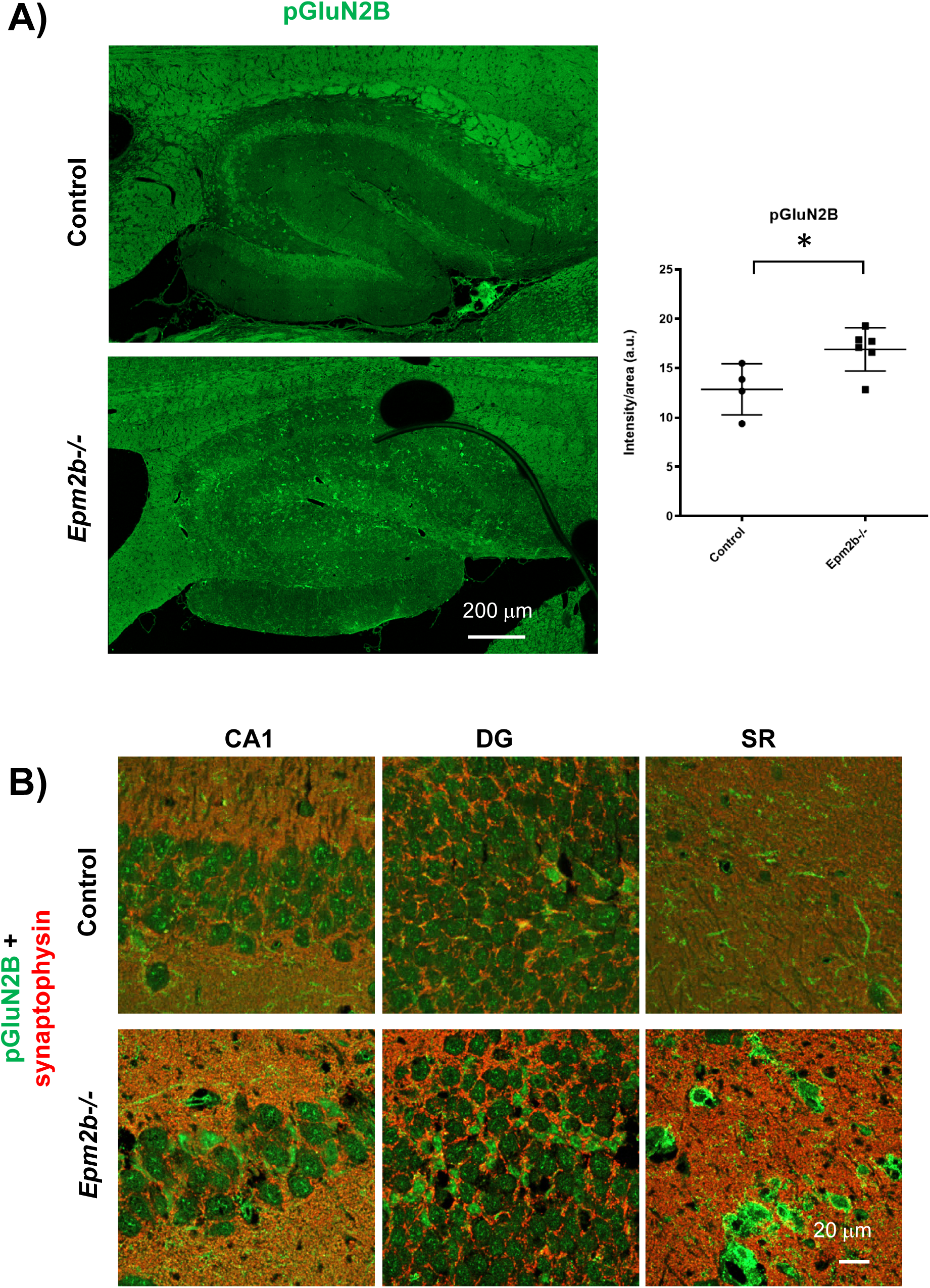
Immunofluorescence analyses of the phosphorylated form of the GluN2B subunit of the NMDA receptor. A) Representative confocal images of the whole hippocampus derived from control and *Epm2b-/-* mice of 16 months of age, labeled by immunofluorescence with anti-pGluN2B antibodies (in green). At least four independent samples from each genotype (males and females) were analyzed in the same way. The scale corresponds to 200 micrometers. The right panel indicates the quantification of the intensity of the corresponding signal in the SR region, related to the analyzed area. Results are expressed as median with a range of at least four independent samples and represented as arbitrary units (a.u.). Differences between the groups were analyzed by Mann-Whitney non-parametric t-test. P-values have been considered as *p<0.05 (Supplementary Table S2). B) A magnification of the Cornu ammonis (CA1), Dentate gyrus (DG), and Stratum radiatum (SR) regions of representative confocal images of the hippocampus derived from control and *Epm2b-/-* mice of 16 months of age, co-labeled by immunofluorescence with anti-pGluN2B (in green) and the pre-synaptic marker anti-synaptophysin (in red) antibodies, is shown. At least four independent samples from each genotype (males and females) were analyzed in the same way. The scale corresponds to 20 micrometers.

**Figure 3:**
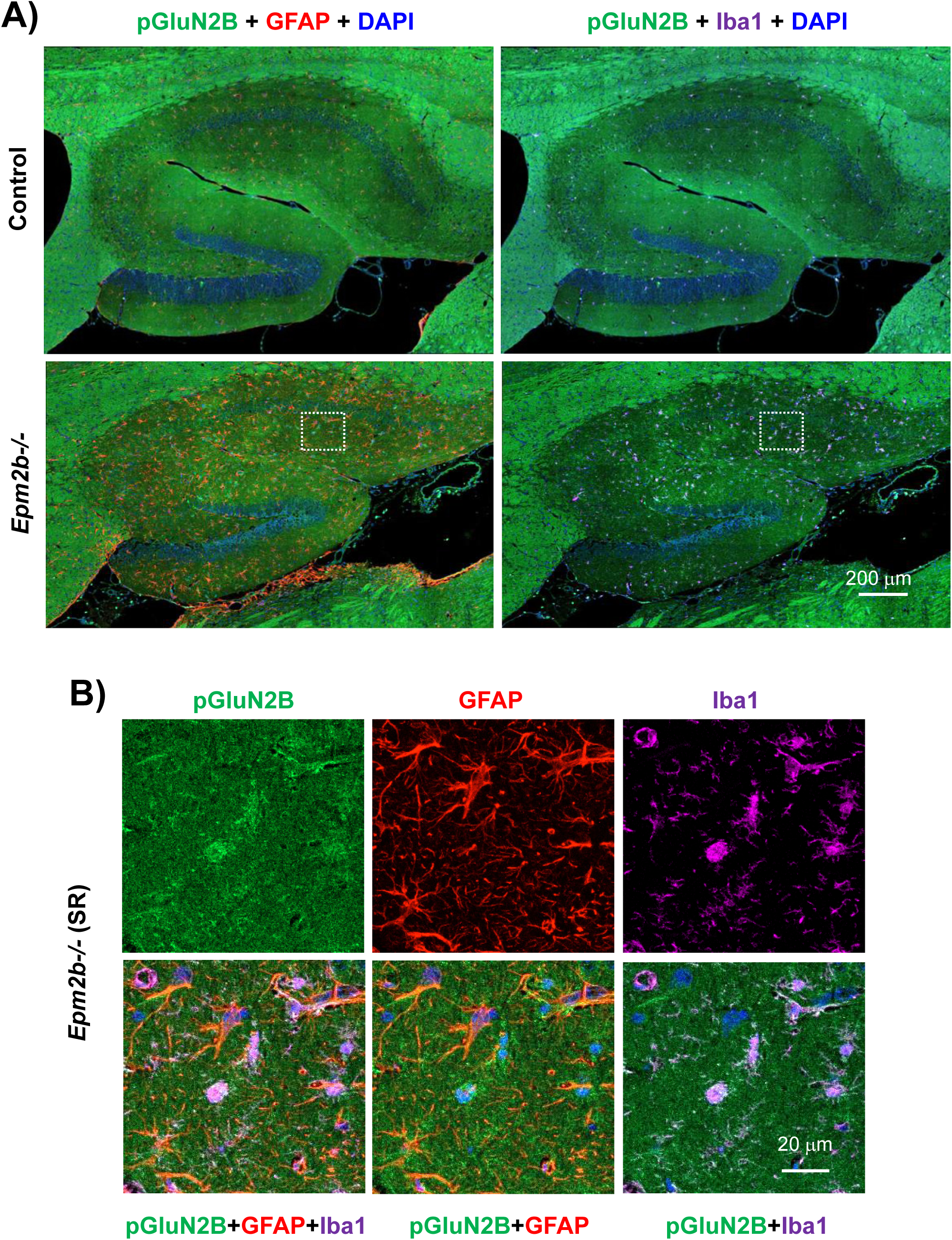
Colocalization of the pGluN2B signal with GFAP and Iba1 markers. A) Similar samples as in Figure 2 were co-labeled with anti-pGluN2B (in green), GFAP (an astrocytic marker; in red), and Iba1 (a microglia marker, in magenta); nuclei were labeled with DAPI (in blue). The scale corresponds to 200 micrometers. B) Different fluorescent channels of a magnification of the SR region of the hippocampus of *Epm2b-/-* samples are shown, corresponding to the dotted square of A). The pGluN2B signal co-localizes with the Iba1 marker. The scale corresponds to 20 micrometers.

#### 1.2. AMPA receptor

Next, we examined the levels of different subunits of the AMPA receptor, namely GluA1, and GluA2. We did not observe major changes in the levels of these subunits between LD and control samples in the hippocampal areas indicated above (Supplementary Fig. S2A and S3A). When we co-labeled the samples with synaptophysin, we did not observe changes in the distribution of these subunits at the CA1 and DG areas between control and LD samples, and the staining of the SR area was also similar between the two groups (Supplementary Fig. S2B and S3B). Western blot analysis did not reveal any difference in the levels of these proteins either (Supplementary Fig. S9A).

Since it has been described that TNF signaling promotes the phosphorylation of different subunits of the AMPA receptor, we analyzed the presence of phosphorylated forms of the GluA2 subunit (pGluA2-S880) alone (Fig. 4A), or in combination with synaptophysin (Fig. 4B). We observed a tendency to an increase of the intensity of the pGluA2 signal in the SR region of the hippocampus of *Epm2b-/-* samples in comparison to controls (25.25 ± 3.02 a.u. in control vs 29.78 ± 7.36 a.u. in *Epm2b-/-*; p-value: 0.111) (Fig. 4A; Supplementary Table S2), but did not observe major changes in the intensity of the neuronal staining of the CA1 and DG areas between control and *Epm2b-/-* samples (not shown). Interestingly, we observed in the CA1 and DG areas of the *Epm2b-/-* samples the presence of ramified cells (white arrow) that were also detected in the SR region (Fig. 4B), which were not present in the control samples. As shown in Fig. 5, the signal of pGluA2 in the SR region of the LD samples corresponded to reactive astrocytes. This assumption is supported by the magnifications presented in Fig. 5B where it is clearly seen that pGluA2 signal co-localizes with reactive astrocytes expressing the GFAP marker and not with activated microglia. Western blot analysis did not show differences between control and LD samples (Supplementary Fig. S9A). In any case, our results indicate that in LD samples astrocytes express the pGluA2 subunit of the AMPA receptor.

**Figure 4:**
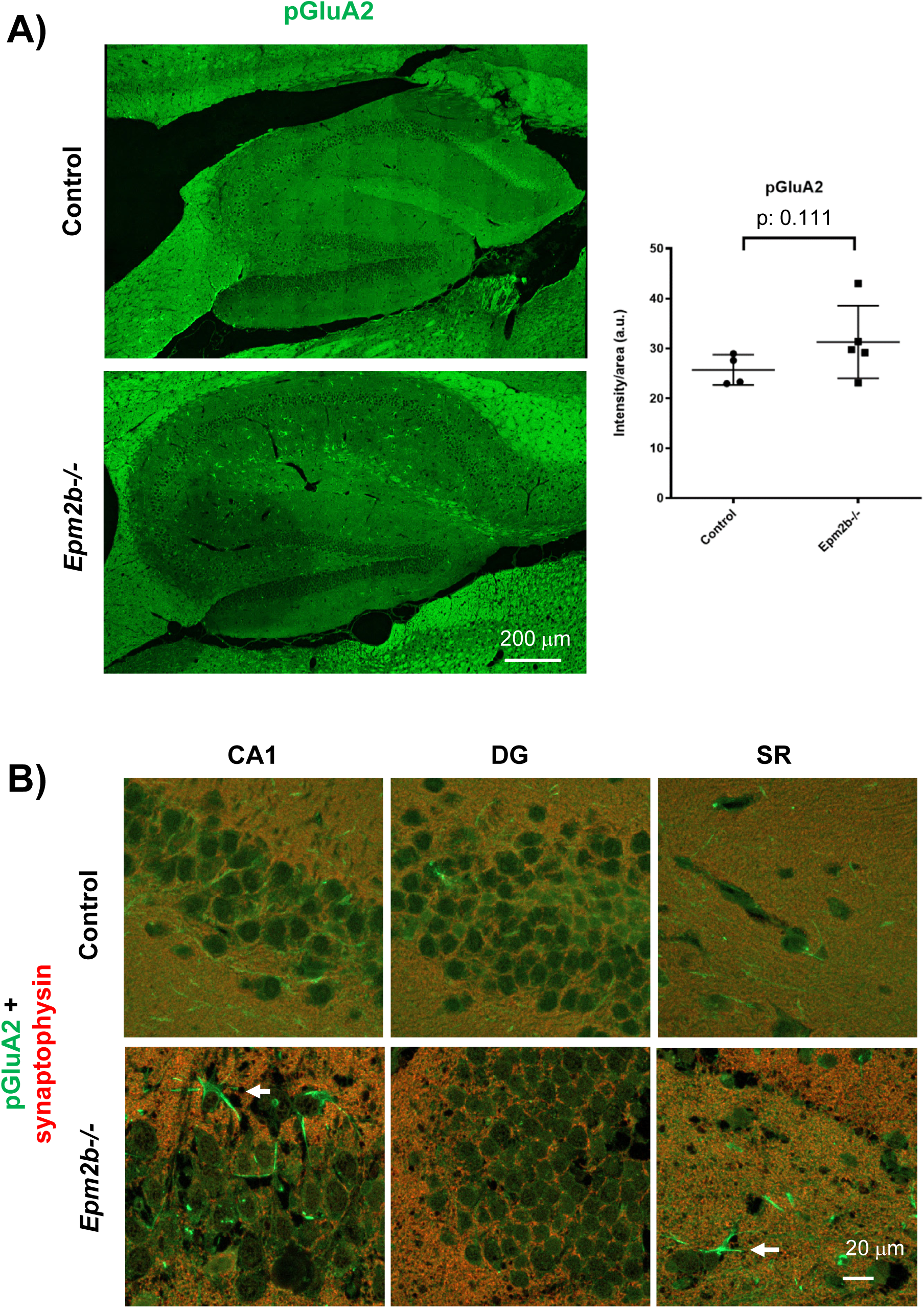
Immunofluorescence analyses of the phosphorylated form of the GluA2 subunit of the AMPA receptor. A) Representative confocal images of the whole hippocampus derived from control and *Epm2b-/-* mice of 16 months of age, labeled by immunofluorescence with anti-pGluA2 antibodies (in green). At least four independent samples from each genotype (males and females) were analyzed in the same way. The scale corresponds to 200 micrometers. The right panel indicates the quantification of the intensity of the corresponding signal in the SR region, related to the analyzed area. Results are expressed as median with a range of at least four independent samples and represented as arbitrary units (a.u.). Differences between the groups were analyzed by Mann-Whitney non-parametric t-test (Supplementary Table S2). B) A magnification of the Cornu ammonis (CA1), Dentate gyrus (DG), and Stratum radiatum (SR) regions of representative confocal images of the hippocampus derived from control and *Epm2b-/-* mice of 16 months of age, co-labeled by immunofluorescence with anti-pGluA2 (in green) and the pre-synaptic marker anti-synaptophysin (in red) antibodies, is shown. White arrows indicate ramified cells. The scale corresponds to 20 micrometers.

**Figure 5:**
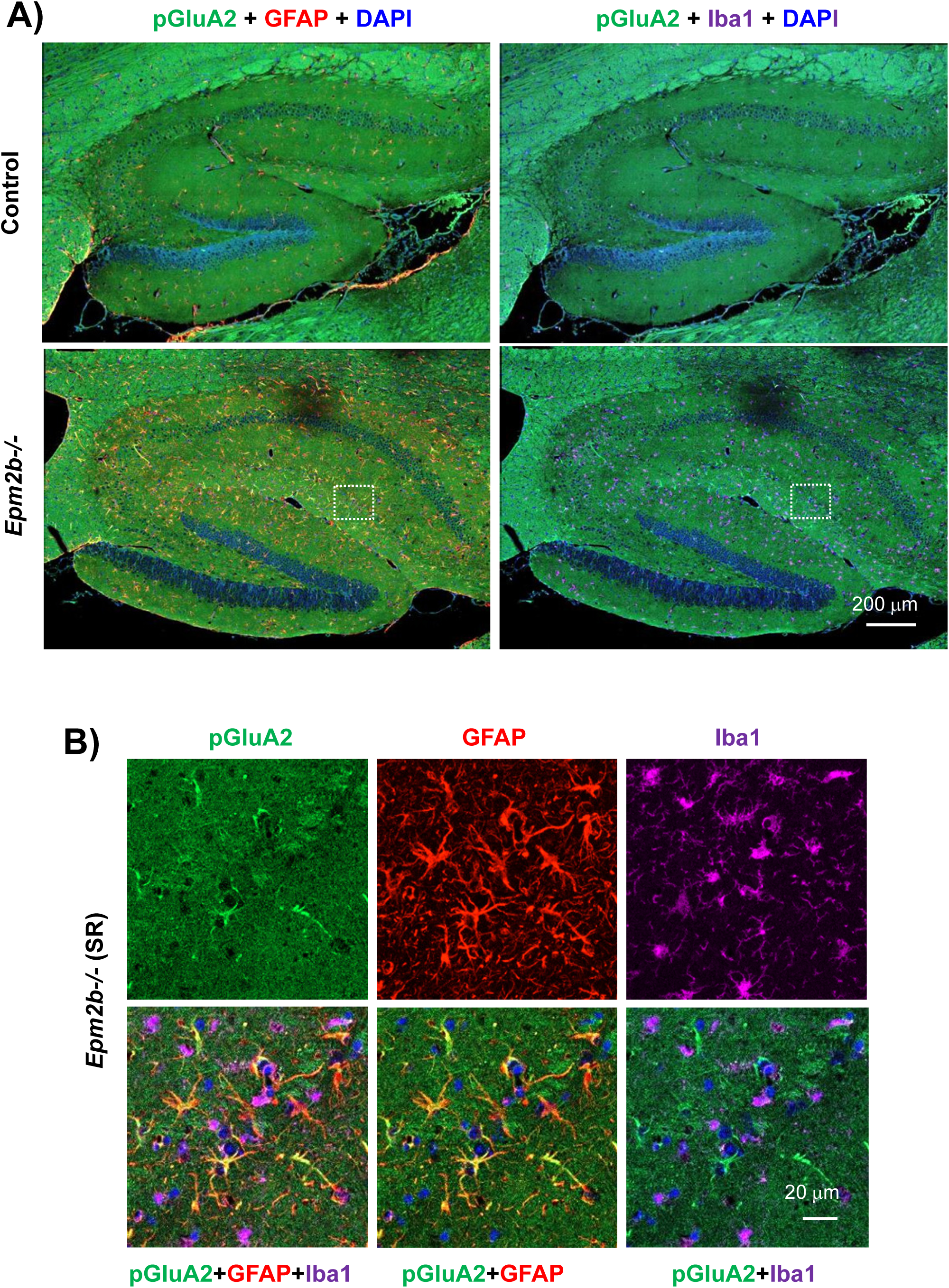
Colocalization of the pGluA2 signal with GFAP and Iba1 markers. A) Similar samples as in Figure 4 were co-labeled with anti-pGluA2 (in green), GFAP (an astrocytic marker; in red), and Iba1 (a microglia marker, in magenta); nuclei were labeled with DAPI (in blue). The scale corresponds to 200 micrometers. B) Different fluorescent channels of a magnification of the SR region of the hippocampus of *Epm2b-/-* samples are shown, corresponding to the dotted square of A). The pGluA2 signal co-localizes with the GFAP marker. The scale corresponds to 20 micrometers.

#### 1.3. Kainate receptor

Then, we analyzed one of the most important subunits of the Kainate glutamate receptor, namely GluK2, alone (Fig. 6A), or in combination with synaptophysin (Fig. 6B). We observed an increase in the intensity of the GluK2 signal in the SR region of the hippocampus of *Epm2b-/-* samples in comparison to controls (11.41 ± 3.91 a.u. in control vs 28.76 ± 16.60 a.u. in *Epm2b-/-* samples; p-value: 0.0173) (Fig. 6A; Supplementary Table S2), but did not observe major changes in the intensity of the neuronal staining of the CA1 and DG areas between control and *Epm2b-/-* samples (not shown). However, at the neuronal level, we observed a punctuated nuclear staining of GluK2 in the LD neurons present in the CA1 and DG areas, which was absent in the controls (Fig. 6B). More importantly, we observed a signal of GluK2 in non-neuronal cells in the CA1 and DG areas of LD samples and in the SR area (Fig. 6B, white arrows). When we performed the colocalization studies of the GluK2 signal with GFAP and Iba1 markers we observed that reactive astrocytes and not activated microglia where the cells which expressed GluK2 in the LD samples (Fig. 7A and 7B). We analyzed the samples by Western blot analysis but, unfortunately, the antibody did not detect any band (Supplementary Fig. S9A) probably because, as the provider indicates in the datasheet, it is specific only for ICC/IF techniques. In any case, our results indicate that in LD samples, astrocytes express the GluK2 subunit of the kainate receptor.

**Figure 6:**
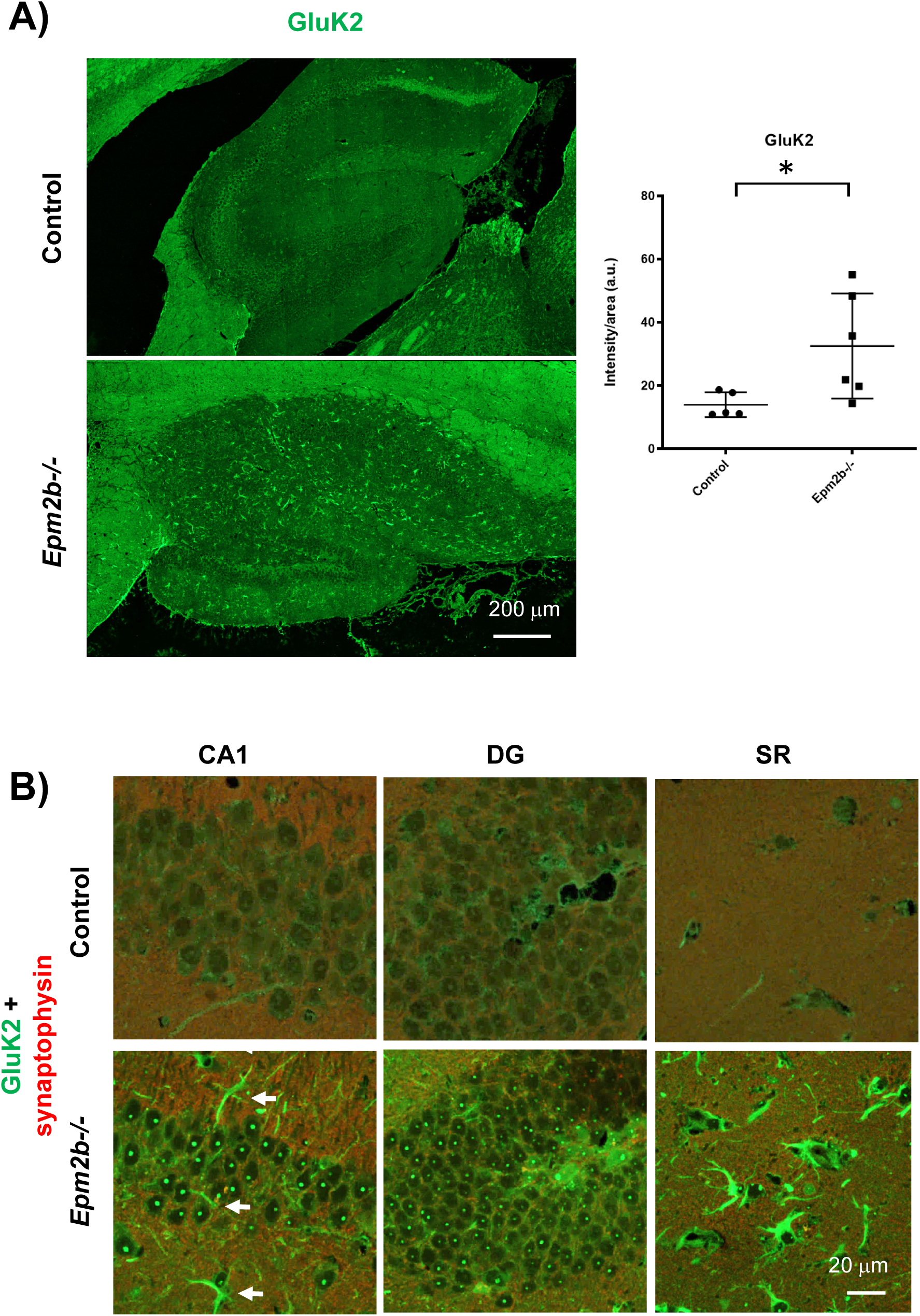
Immunofluorescence analyses of the GluK2 subunit of the Kainate receptor. A) Representative confocal images of the whole hippocampus derived from control and *Epm2b-/-* mice of 16 months of age, labeled by immunofluorescence with anti-GluK2 antibodies (in green). At least four independent samples from each genotype (males and females) were analyzed in the same way. The scale corresponds to 200 micrometers. The right panel indicates the quantification of the intensity of the corresponding signal in the SR region, related to the analyzed area. Results are expressed as median with a range of at least four independent samples and represented as arbitrary units (a.u.). Differences between the groups were analyzed by Mann-Whitney non-parametric t-test. P-values have been considered as *p<0.05 (Supplementary Table S2). B) A magnification of the Cornu ammonis (CA1), Dentate gyrus (DG), and Stratum radiatum (SR) regions of representative confocal images of the hippocampus derived from control and *Epm2b-/-* mice of 16 months of age, co-labeled by immunofluorescence with anti-GluK2 (in green) and the pre-synaptic marker anti-synaptophysin (in red) antibodies, is shown. White arrows indicate ramified cells. The scale corresponds to 20 micrometers.

**Figure 7:**
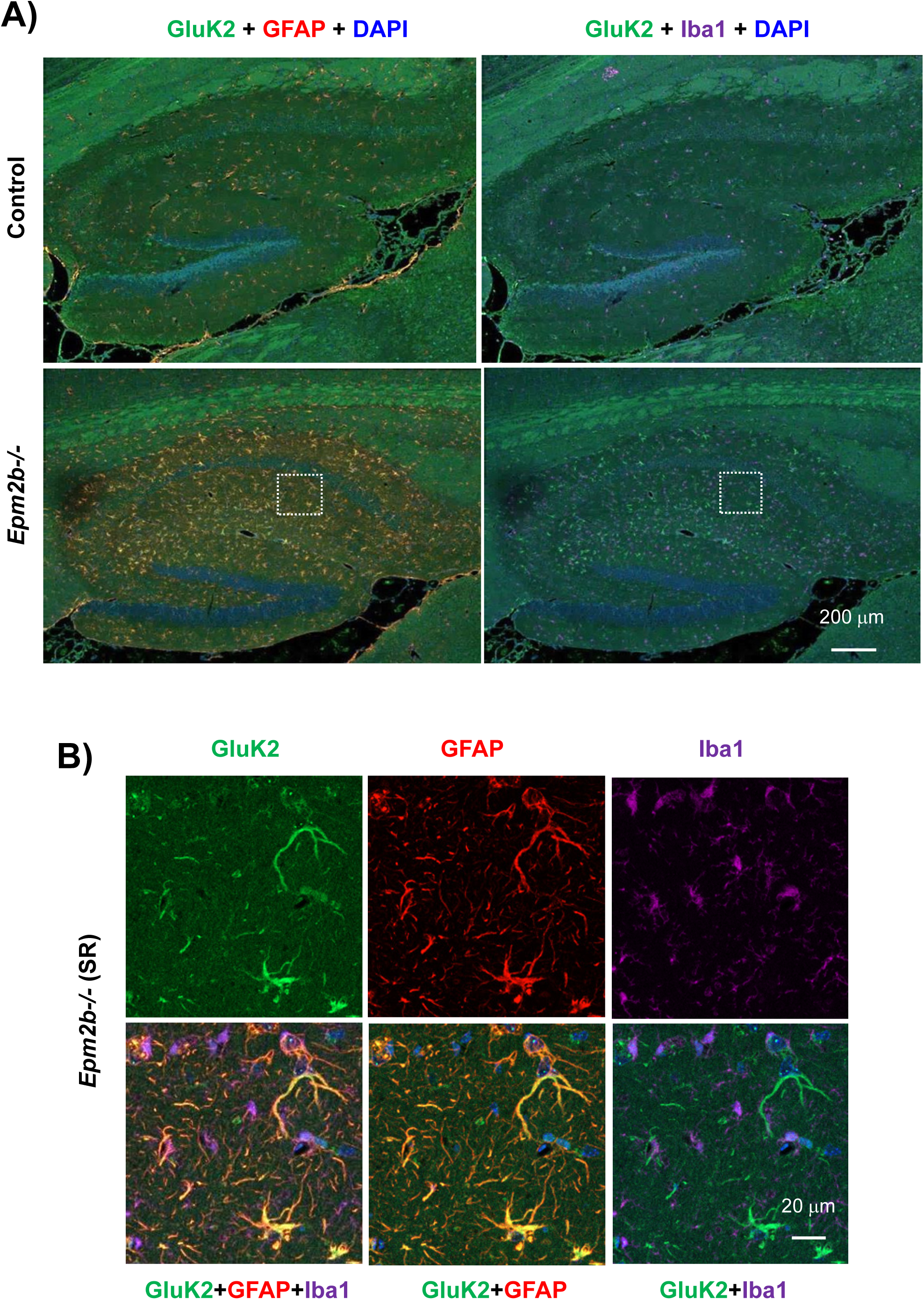
Colocalization of the GluK2 signal with GFAP and Iba1 markers. A) Similar samples as in Figure 6 were co-labeled with anti-GluK2 (in green), GFAP (an astrocytic marker; in red), and Iba1 (a microglia marker, in magenta); nuclei were labeled with DAPI (in blue). The scale corresponds to 200 micrometers. B) Different fluorescent channels of a magnification of the SR region of the hippocampus of *Epm2b-/-* samples are shown, corresponding to the dotted square in A). The GluK2 signal co-localizes with the GFAP marker. The scale corresponds to 20 micrometers.

#### 1.4. Metabotropic Glutamate receptor

Finally, we analyzed the levels of the mGluR5 isoform of the metabotropic glutamate receptor. However, in the hippocampus, we did not observe major differences in the levels of this isoform in samples from LD animals in comparison to controls, either by immunofluorescence (Supplementary Fig. S4A), combined immunofluorescence with synaptophysin (Supplementary Fig. S4B), or by Western blot analysis (Supplementary Fig. S9A).

### 2. GABAergic signaling

Since it has been reported that TNF also diminishes GABAergic signaling, we analyzed the levels of different proteins related to this pathway.

#### 2.1. GABAA receptor

We analyzed the levels of GABAA-α1 and GABAA-γ2 subunits of the ionotropic GABAA receptor. In both cases, we did not observe major changes in the levels of these subunits between LD and control samples by immunofluorescence (Supplementary Fig. S5A and S6A), combined immunofluorescence with synaptophysin (Supplementary Fig. S5B for GABAA-α1), or by Western blot analysis (Supplementary Fig. S9B). We also analyzed the levels of the GAD65/67 marker, related to GABAergic neurons, and we did not find any differences either (Supplementary Fig. S6B and Fig. S9B).

#### 2.2. GABA transporters

Then, we analyzed the levels of the two main GABA transporters, namely GAT1 and GAT3. In the case of GAT1, we observed an increase in the intensity of the GAT1 signal in the SR region of the hippocampus of *Epm2b-/-* samples in comparison to controls (16.45 ± 4.50 a.u. in control vs 24.78 ± 12.81 a.u. in *Epm2b-/-* samples; p-value: 0.0260) (Fig. 8A; Supplementary Table S2). No major differences were observed in the intensity of the neuronal staining of the CA1 and DG areas (not shown), but in the case of the SR region of LD samples, we observed the presence of more abundant non-neuronal cells that were labeled with the anti-GAT1 antibody (Fig. 8B). These cells corresponded to reactive astrocytes and not to microglia, as demonstrated in the co-localization experiments indicated in Fig. 9A and 9B. Western blot analysis did not show differences between control and LD samples (Supplementary Fig. S9B). In any case, our results indicate that reactive astrocytes of LD samples express the GAT1 protein.

**Figure 8:**
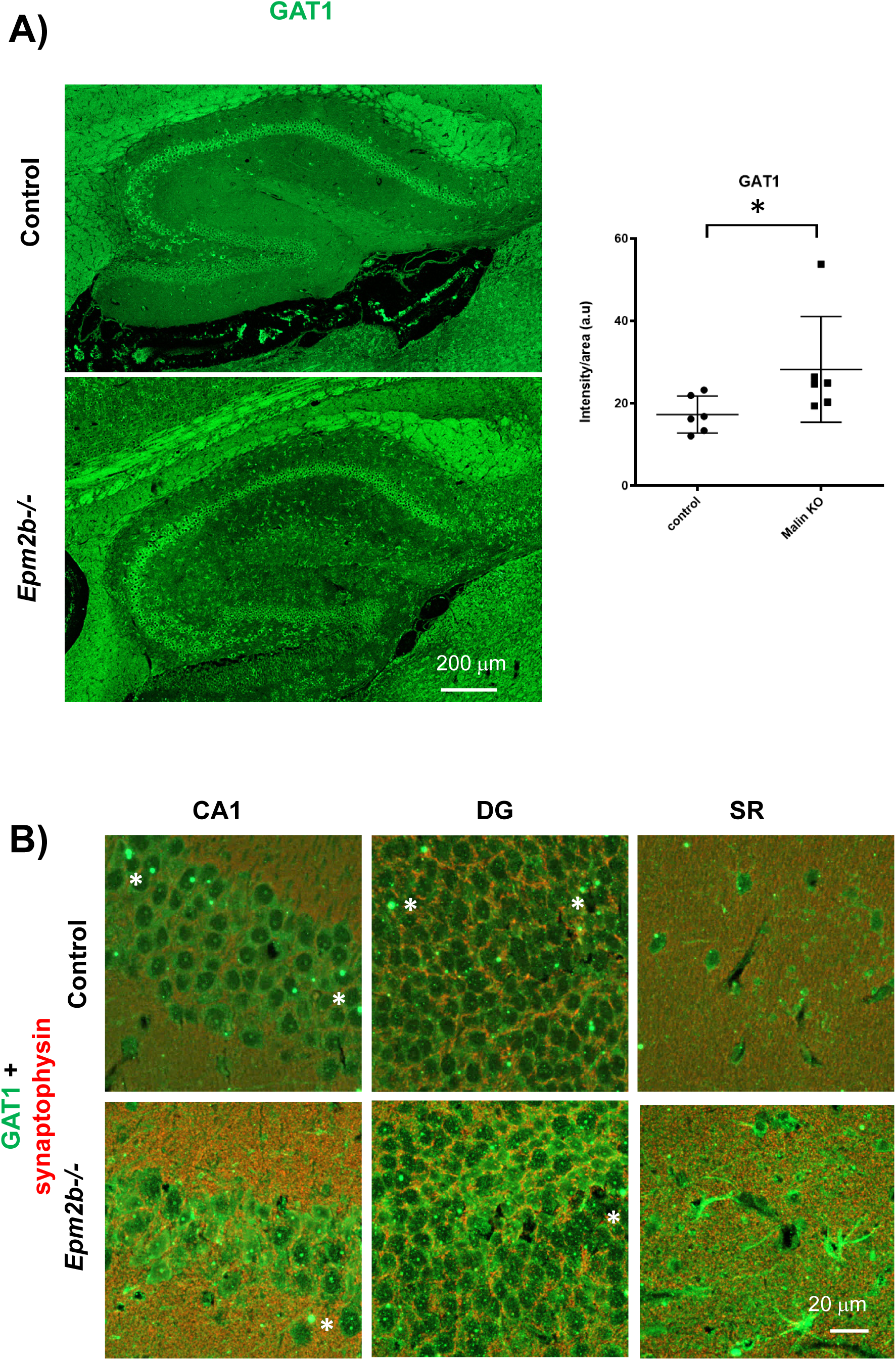
Immunofluorescence analyses of the GAT1 transporter. A) Representative confocal images of the whole hippocampus derived from control and *Epm2b-/-* mice of 16 months of age, labeled by immunofluorescence with anti-GAT1 antibodies (in green). At least four independent samples from each genotype (males and females) were analyzed in the same way. The scale corresponds to 200 micrometers. The right panel indicates the quantification of the intensity of the corresponding signal in the SR region, related to the analyzed area. Results are expressed as median with a range of at least four independent samples and represented as arbitrary units (a.u.). Differences between the groups were analyzed by Mann-Whitney non-parametric t-test. P-values have been considered as *p<0.05 (Supplementary Table S2). B) A magnification of the Cornu ammonis (CA1), Dentate gyrus (DG), and Stratum radiatum (SR) regions of representative confocal images of the hippocampus derived from control and *Epm2b-/-* mice of 16 months of age, co-labeled by immunofluorescence with anti-GAT1 (in green) and the pre-synaptic marker anti-synaptophysin (in red) antibodies, is shown. The scale corresponds to 20 micrometers. The antibody produced artefactual punctuated staining (asterisks) in both control and *Epm2b-/-* samples.

**Figure 9:**
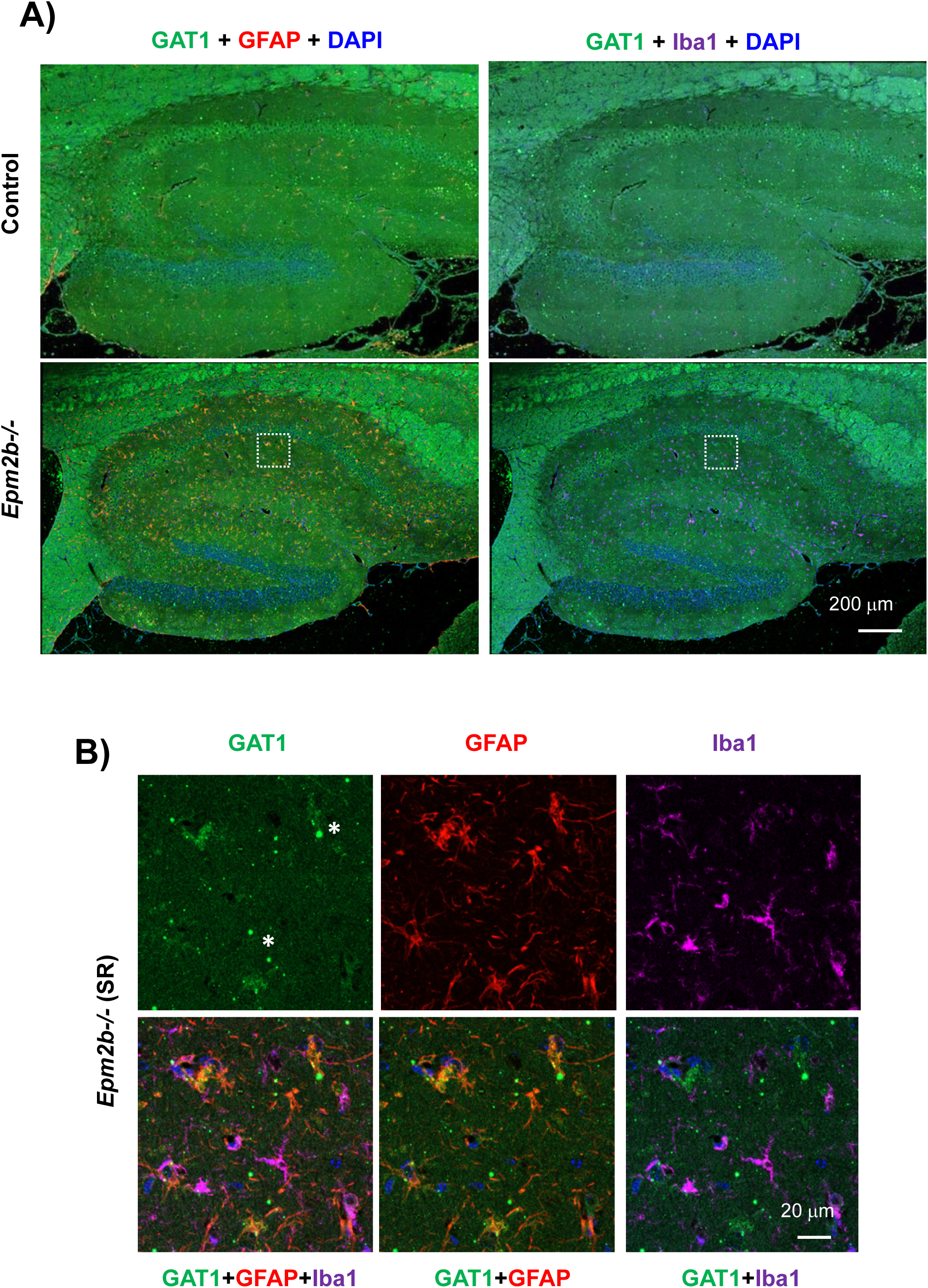
Colocalization of the GAT1 signal with GFAP and Iba1 markers. A) Similar samples as in Figure 8 were co-labeled with anti-GAT1 (in green), GFAP (an astrocytic marker; in red), and Iba1 (a microglia marker, in magenta); nuclei were labeled with DAPI (in blue). The scale corresponds to 200 micrometers. B) Different fluorescent channels of a magnification of the SR region of the hippocampus of *Epm2b-/-* samples are shown, corresponding to the dotted square in A). The GAT1 signal co-localizes with the GFAP marker. The scale corresponds to 20 micrometers. The antibody produced artefactual punctuated staining (asterisks) in both control and *Epm2b-/-* samples.

In the case of the GAT3 protein, we did not observe major differences in the levels of this isoform in samples from LD animals in comparison to controls by immunofluorescence (Supplementary Fig. S7A), combined immunofluorescence with synaptophysin (Supplementary Fig. S7B), or by Western blot analysis (Supplementary Fig. 9B).

### 3. Src kinase family

TNF signaling leads to the activation of different protein kinases [15], [41]. Among them, one important group of kinases related to neurotransmitter signaling is the Src family of protein kinases. Since we observed changes in the levels of phosphorylated forms of GluN2B and GluA2 subunits, we decided to study the presence of activated forms of three members of this kinase family, namely Src, Lyn, and Fyn.

#### 3.1. Src kinase

We analyzed the levels of Src kinase but, at the hippocampal levels, did not observe major differences between LD and control samples by immunofluorescence or Western blot analysis (Supplementary Fig. S8 and Fig. S9C, respectively). However, when we analyzed by immunofluorescence the levels of active phosphorylated forms of Src kinase (pSrc-Y416), we detected an increase in the levels of this form in the SR region of the hippocampus of LD samples in comparison to controls (22.38 ± 2.23 a.u. in control vs 32.94 ± 4.18 a.u. in *Epm2b-/-* samples; p-value: 0.0159) (Fig. 10A; Supplementary Table S2), but did not observe major changes in the intensity of the neuronal staining of the CA1 and DG areas between control and *Epm2b-/-* samples (not shown). Interestingly, we observed the appearance of non-neuronal cells that were pSrc+ in the CA1 and SR regions (Fig. 10B, white arrows). This non-neuronal signal co-localized mainly with activated microglia, as indicated by the colocalization studies of the pSrc signal and the GFAP and Iba1 markers shown in Fig. 11A and 11B. Western blot analysis did not show differences between control and LD samples (Supplementary Fig. S9C). In any case, our results indicate that, in LD samples, microglial cells express the phosphorylated form of Src kinase.

**Figure 10:**
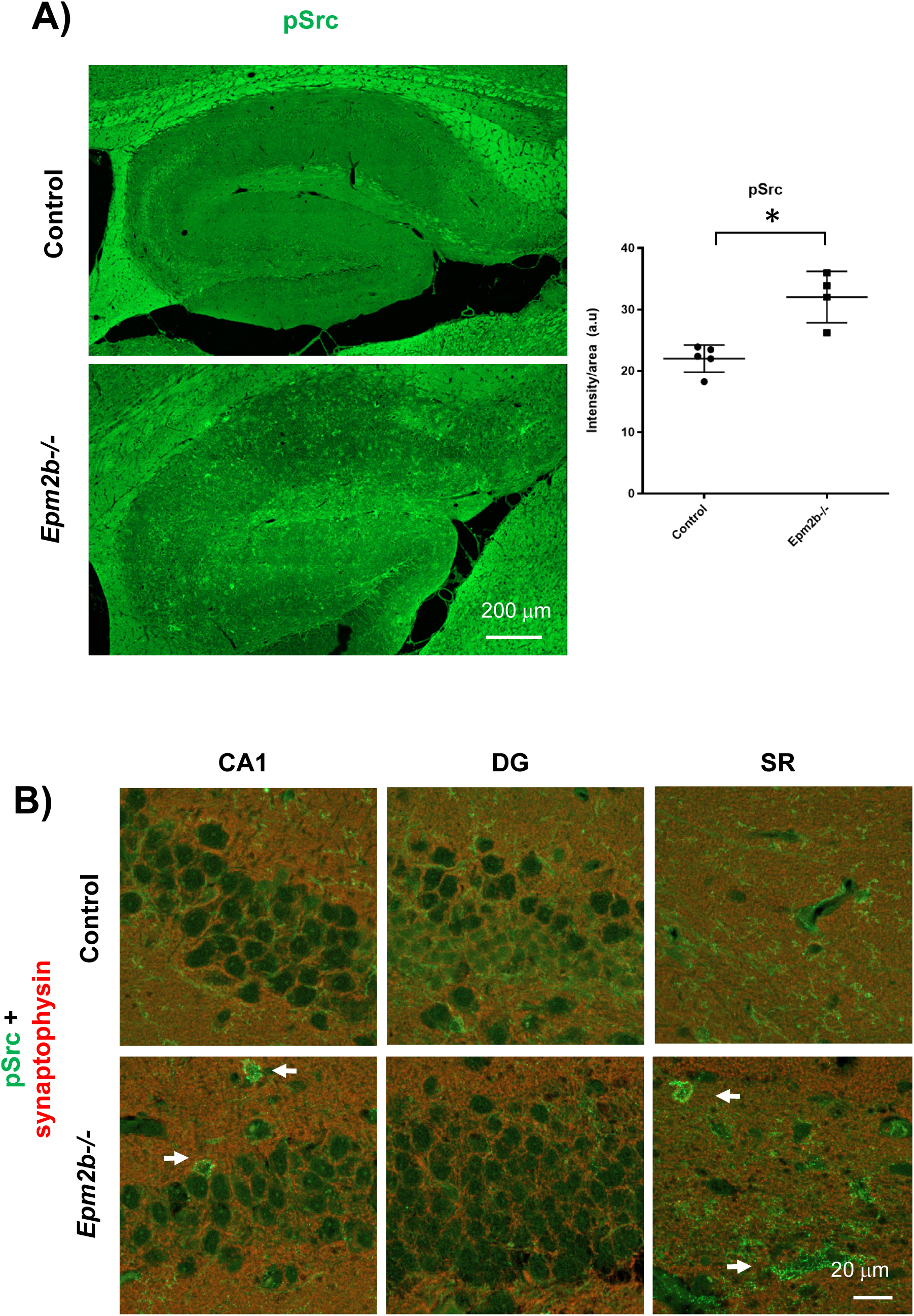
Immunofluorescence analyses of the phosphorylated form of the Src protein kinase. A) Representative confocal images of the whole hippocampus derived from control and *Epm2b-/-* mice of 16 months of age, labeled by immunofluorescence with anti-pSrc antibodies (in green). At least four independent samples from each genotype (males and females) were analyzed in the same way. The scale corresponds to 200 micrometers. The right panel indicates the quantification of the intensity of the corresponding signal in the SR region, related to the analyzed area. Results are expressed as median with a range of at least four independent samples and represented as arbitrary units (a.u.). Differences between the groups were analyzed by Mann-Whitney non-parametric t-test. P-values have been considered as *p<0.05 (Supplementary Table S2). B) A magnification of the Cornu ammonis (CA1), Dentate gyrus (DG), and Stratum radiatum (SR) regions of representative confocal images of the hippocampus derived from control and *Epm2b-/-* mice of 16 months of age, co-labeled by immunofluorescence with anti-pSrc (in green) and the pre-synaptic marker anti-synaptophysin (in red) antibodies, is shown. White arrows indicate rounded cells. The scale corresponds to 20 micrometers.

**Figure 11:**
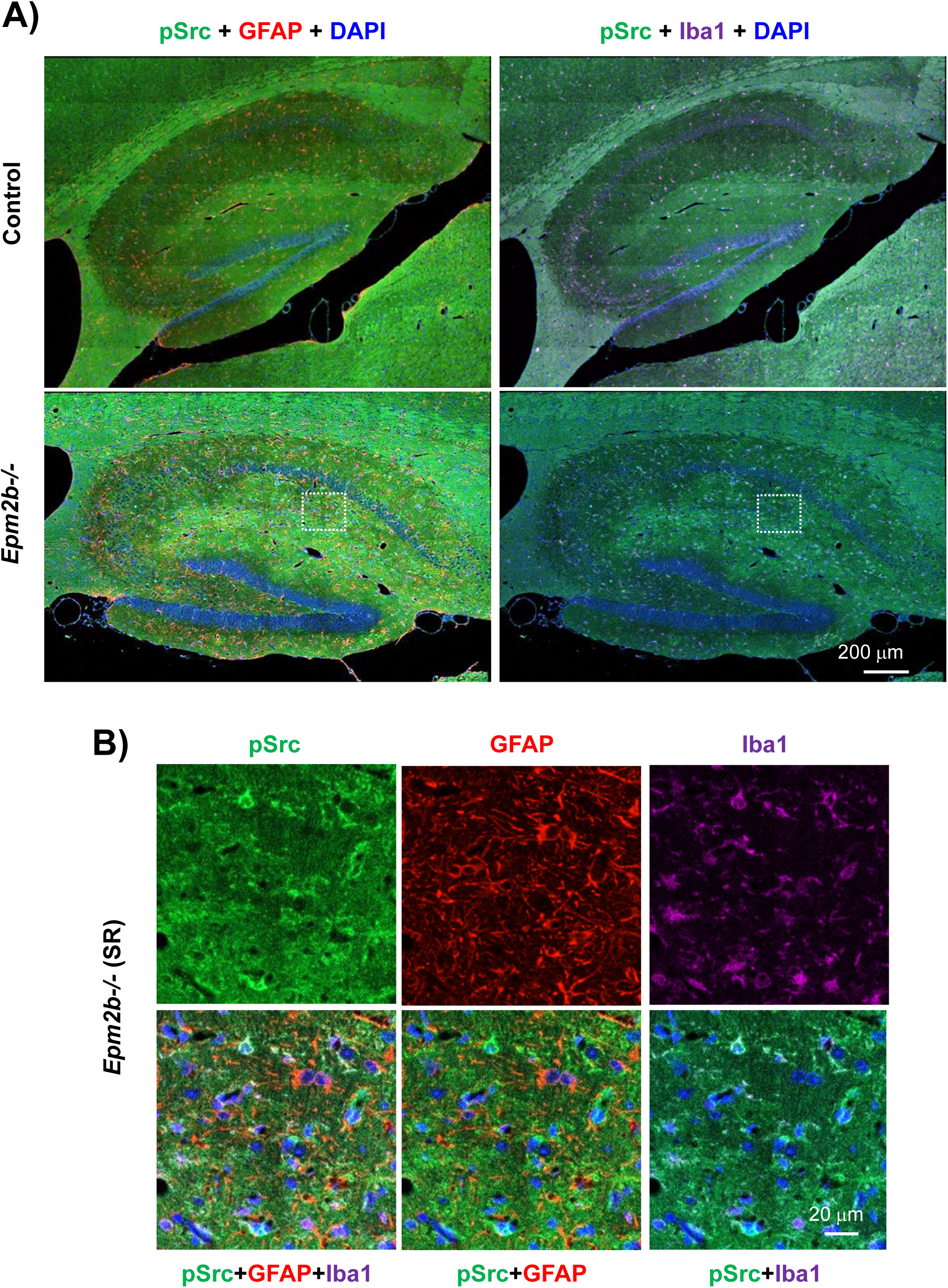
Colocalization of the pSrc signal with GFAP and Iba1 markers. A) Similar samples as in Figure 8 were co-labeled with anti-pSrc (in green), GFAP (an astrocytic marker; in red), and Iba1 (a microglia marker, in magenta); nuclei were labeled with DAPI (in blue). The scale corresponds to 200 micrometers. B) Different fluorescent channels of a magnification of the SR region of the hippocampus of *Epm2b-/-* samples are shown, corresponding to the dotted square in A). The pSrc signal co-localizes with Iba1 marker. The scale corresponds to 20 micrometers.

#### 3.2. Lyn kinase

The production of this kinase followed the same trend as the one from the Src kinase defined above. No major differences were observed in the hippocampus of LD animals in comparison to control by immunofluorescence or Western blot analysis (Supplementary Fig. S8 and Fig. S9C). However, when we analyzed by immunofluorescence the levels of active phosphorylated forms of Lyn kinase (pLyn-Y397), we observed an increase in the signal of this kinase in the SR region of the hippocampus of LD animals in comparison to controls (18.64 ± 3.60 in control vs 30.49 ± 6.40 a.u. in *Epm2b-/-* samples; p-value: 0.0317) (Fig. 12A; Supplementary Table S2), but did not observe major changes in the intensity of the neuronal staining of the CA1 and DG areas between control and *Epm2b-/-* samples (not shown). We also observed the appearance of non-neuronal cells that were pLyn+ (Fig. 12B; ramified cells in the CA1 and SR region, white arrows). This staining co-localized mainly with reactive astrocytes as indicated by the co-localization experiments of the pLyn signal with the GFAP and Iba1 markers shown in Fig. 13A and 13B. Western blot analysis did not show differences between control and LD samples (Supplementary Fig. S9C). In any case, our results indicate that, in LD samples, astrocytes express the phosphorylated form of Lyn kinase

**Figure 12:**
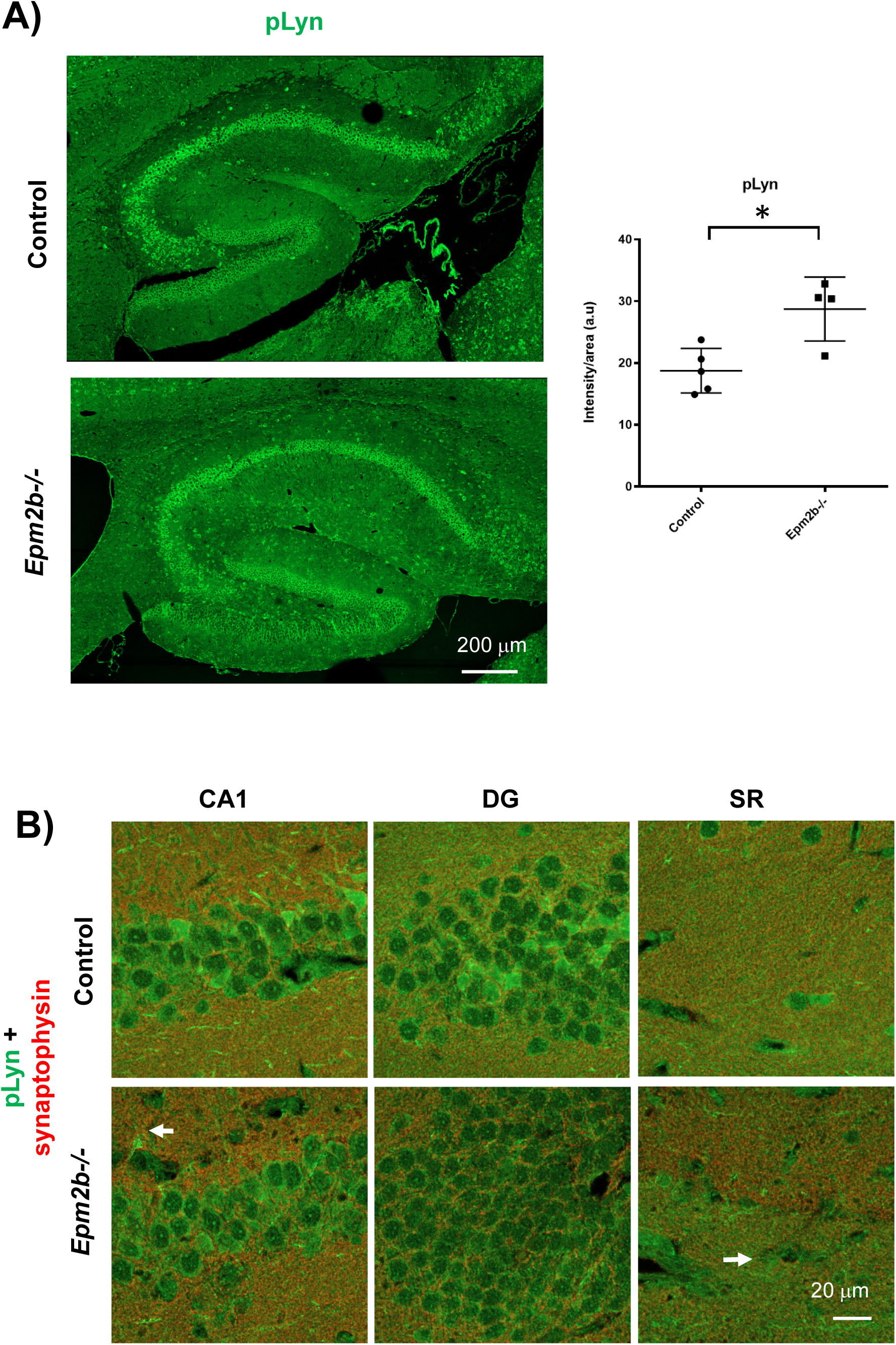
Immunofluorescence analyses of the phosphorylated form of the Lyn protein kinase. A) Representative confocal images of the whole hippocampus derived from control and *Epm2b-/-* mice of 16 months of age, labeled by immunofluorescence with anti-pLyn antibodies (in green). At least four independent samples from each genotype (males and females) were analyzed in the same way. The scale corresponds to 200 micrometers. The right panel indicates the quantification of the intensity of the corresponding signal in the SR region, related to the analyzed area. Results are expressed as median with a range of at least four independent samples and represented as arbitrary units (a.u.). Differences between the groups were analyzed by Mann-Whitney non-parametric t-test. P-values have been considered as *p<0.05 (Supplementary Table S2). B) A magnification of the Cornu ammonis (CA1), Dentate gyrus (DG), and Stratum radiatum (SR) regions of representative confocal images of the hippocampus derived from control and *Epm2b-/-* mice of 16 months of age, co-labeled by immunofluorescence with anti-pLyn (in green) and the pre-synaptic marker anti-synaptophysin (in red) antibodies, is shown. The scale corresponds to 20 micrometers.

**Figure 13:**
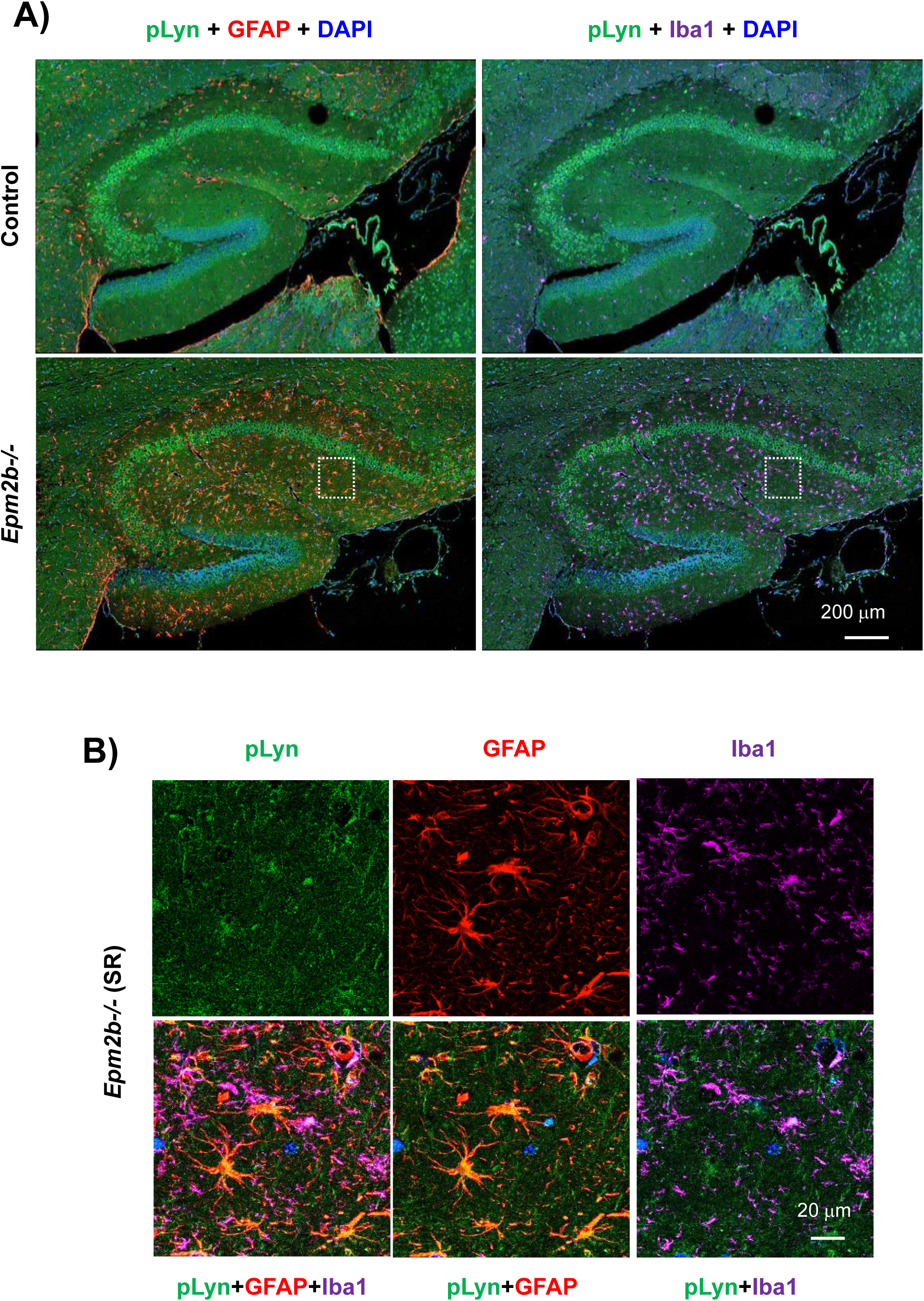
Colocalization of the pLyn signal with GFAP and Iba1 markers. A) Similar samples as in Figure 12 were co-labeled with anti-pLyn (in green), GFAP (an astrocytic marker; in red), and Iba1 (a microglia marker, in magenta); nuclei were labeled with DAPI (in blue). The scale corresponds to 200 micrometers. B) Different fluorescent channels of a magnification of the SR region of the hippocampus of *Epm2b-/-* samples are shown, corresponding to the dotted square in A). The pLyn signal co-localizes with the GFAP marker. The scale corresponds to 20 micrometers.

#### 3.3. Fyn kinase

Finally, we analyzed the levels of Fyn kinase and also active phosphorylated forms of this kinase (pFyn-Y530) but in both cases, we did not observe major changes between LD and control samples by immunofluorescence (Supplementary Fig. S8) or Western blot analysis (Supplementary Fig. S9C).

## DISCUSSION

Lafora disease (LD) is characterized by progressive myoclonus epilepsy. Previous work of our group demonstrated that the uptake of glutamate is impaired in the astrocytes from LD mouse models [13], [42], and the brain of LD animals contains higher levels of the excitatory glutamate neurotransmitter [43]. In agreement with this, it has been recently demonstrated that the brains of Lafora disease patients contain higher levels of glutamate/glutamine [44]. In addition, recent work indicates that neuroinflammation should be considered a new trait in LD [14]. In animal models of LD, we demonstrated the presence of reactive astrocytes, activated microglia, and infiltrated T-lymphocytes in the brain, which activated TNF and IL-6 signaling pathways leading to the overexpression of different pro-inflammatory mediators, e.g., NF-kB, CXCL10, and Lcn2 [14], [15].

Since the activation of the TNF signaling pathway contributes to neuronal hyper-excitability [16], in this work we present evidence that in an animal model of LD (*Epm2b-/-* mice) there is an upregulation of different subunits of the three major excitatory glutamate receptors, namely NMDA, AMPA, and Kainate receptors. Although by RNAseq analysis performed in whole brain, we did not find changes in the expression of the corresponding genes when comparing controls vs LD samples [14], in this work we found differences in the levels of several subunits of these receptors only when we analyzed by immunofluorescence the Stratum radiatum area of the hippocampus. No differences in these subunits were observed in other analyzed areas (e.g., Cornu ammonis CA1, CA3, or DG). The specific expression of these subunits in one particular area of the hippocampus is perhaps the reason why we could not detect differences in the levels of these proteins by Western blot analysis since samples prepared for this technique contained the whole hippocampus.

In the case of NMDA receptors, we show that the phosphorylated form of the GluN2B subunit (at residue Y1336) is overrepresented in the hippocampus of LD animals in comparison to controls and that this overexpression corresponds to an increase in the presence of this subunit in activated microglia. The fact that we found no differences in the levels of un-phosphorylated forms of this subunit may indicate that GluN2B gets activated in the hippocampus of the *Epm2b-/-* samples. The higher detection of the pGluN2B signal at the hippocampal level could be due to a higher number of activated microglia expressing this form and/or to a higher expression of the subunit in these cells. In any case, the huge production of pGluN2B at the hippocampal level may affect the overall response to glutamate and this may contribute to hyper-excitability [32]. It is important to note that the Cornu ammonis 1 (CA1) region of the hippocampus is especially sensitive to the activation of NMDA receptors [45]. It is also worth pointing out that in temporal lobe epilepsy (TLE), there is an increase of NMDA receptor subunits in the hippocampus of affected patients [20], [23], connecting in this way activation of NMDA receptors and pathology.

In the case of AMPA receptors, we observed a tendency to increase levels of phosphorylated forms of the GluA2 subunit (at residue S880) in the hippocampus of LD animals. The pGluA2 signal corresponded to the production of this subunit by reactive astrocytes. Since we found no differences in the levels of un-phosphorylated forms of this subunit, this may indicate that GluA2 gets activated in the hippocampus of the *Epm2b-/-* samples. Again, the detection of the pGluA2 signal at the hippocampal level could be due to a higher number of reactive astrocytes expressing this form and/or to a higher expression of the subunit in these cells. The activation of AMPA receptors in astrocytes could lead to an activation of NMDA receptors aggravating excitotoxicity [20], [23], [46]. In fact, in patients with status epilepticus, the activity of the AMPA receptors is enhanced and this leads to an increase in the activity of the NMDA receptors, leading to seizures [23].

In the case of Kainate receptors, we observed an increase in the levels of the GluK2 subunit in the hippocampus of LD animals. Receptors containing the GluK2 subunit have been linked to epilepsy, and it has been proposed that its expression in the CA3 pyramidal neurons may cause its neurodegeneration [30], [47], [48]. In LD samples we observed a different subcellular distribution of the GluK2 signal in neurons from CA1 and DG, with a clear punctuated nuclear staining, but at this time we do not know the physiological meaning of this observation. Reactive astrocytes, but not activated microglia, were able to produce the GluK2 subunit. This is interesting since it has been reported that only astrocytes from mice under status epilepticus conditions express in their hippocampus different subunits of Kainate receptors (GluK1, 2, 4 and 5) [49]. These authors indicated that astrocytes from the CA1, CA3, and entorhinal cortex co-expressed Kainate receptor subunits and GFAP markers, but not astrocytes from other brain areas. Therefore, our finding of the presence of the GluK2 subunit in astrocytes from the hippocampus reinforces the link between the expression of GluKRs in reactive astrocytes and epilepsy [46], [48], [49].

We also present evidence that members of the Src family of protein kinases are activated in the hippocampus of LD animals. The presence of active phosphorylated forms of Src (at residue Y416) and Lyn (at residue Y397) kinases may participate in the phosphorylation of different glutamatergic receptor subunits (possibly GluN2B and GluA2, see above), enhancing in this way their activity. We also found that pSrc-Y416 is mainly expressed in activated microglia, whereas pLyn-Y397 is expressed mainly in reactive astrocytes. This may suggest a particular action of these two kinases in each particular cell host.

On the other hand, we also present evidence of alterations in the GABAergic signaling. In this case, we did not find changes in the levels of different GABAergic receptor subunits but in the levels of the GAT1 transporter. We found an increase in the levels of GAT1 in the hippocampus of LD animals, mainly at the level of reactive astrocytes. The higher levels of GAT1 transporter may decrease the levels of GABA in the brain parenchyma, decreasing in this way the inhibitory tone, leading to hyper-excitability [35], [50]. In agreement with this, a recent report indicates that the brain of Lafora disease patients contains lower levels of GABA neurotransmitter [44].

In summary, we present evidence that, in the hippocampus of the LD model used in this work (*Epm2b-/-* mice), there is a dysregulation in the levels of different subunits of the ionotropic glutamatergic receptors in non-neuronal cells such as activated microglia (pGluN2B) and reactive astrocytes (pGluA2 and GluK2), which could increase Ca++ signaling and facilitate the hyperexcitability present in this model [46]. We also show an increase in the levels of the GABA transporter GAT1 in reactive astrocytes, which could diminish the available levels of synaptic GABA, decreasing in this way the inhibitory tone. In addition, we also present evidence of higher levels of active phosphorylated forms of members of the Src protein kinase family in activated microglia (pSrc) and reactive astrocytes (pLyn), which could be responsible for the changes in the activity of the glutamatergic subunits indicated above. All these changes could explain the enhanced sensitivity of LD mice to the effects of pro-convulsant drugs such as kainate or pentylenetetrazole (PTZ) [12] and also the beneficial effect that drugs inhibiting NMDA signaling (e.g. memantine) or AMPA signaling (e.g. perampanel) have in mouse models and patients of LD [51], [52], [53]. Finally, our results highlight the role of reactive astrocytes and activated microglia as key players in LD pathophysiology and define them as therapeutic targets of intervention for finding a treatment for this devastating disease.

## ACKNOWLEDGMENTS

We thank Dr. María Adelaida García-Gimeno for the critical reading of the manuscript.

## DECLARATIONS

## Ethics approval

This study was carried out in strict accordance with the recommendations in the Guide for the Care and Use of Laboratory Animals of the Consejo Superior de Investigaciones Científicas (CSIC, Spain) and approved by the Consellería de Agricultura, Medio Ambiente, Cambio Climatico y Desarrollo Rural from The Generalitat Valenciana. All procedures were approved by the animal committee of the Instituto de Biomedicina de Valencia, CSIC (authorization number IBV-51). All efforts were made to minimize animal suffering. When planning the experiments, the principles outlined in the ARRIVE guidelines and the Basel declaration including the 3R concept have been considered.

## Consent to participate

Not applicable

## Consent for publication

Not applicable

## Availability of data and materials

All data generated or analyzed during this study are included in this published article and its supplementary information files.

## Code availability

Not applicable

## Author’s contribution

RV, TR, and AC-R performed the experiments. TR and PS analyzed the data. PS wrote the manuscript. All authors have read and approved the final version of the manuscript.

## REFERENCES

1. Lafora GR, Glueck B (1911) Beitrag zur histogpathologie der myoklonischen epilepsie. Gesamte Neurol Psychiatr 6:1–14.

2. Garcia-Gimeno MA, Knecht E, Sanz P (2018) Lafora Disease: A Ubiquitination-Related Pathology. Cells 7 (8):87.

3. Pondrelli F, Muccioli L, Licchetta L, Mostacci B, Zenesini C, Tinuper P, Vignatelli L, Bisulli F (2021) Natural history of Lafora disease: a prognostic systematic review and individual participant data meta-analysis. Orphanet J Rare Dis 16 (1):362.

4. Minassian BA, Lee JR, Herbrick JA, Huizenga J, Soder S, Mungall AJ, Dunham I, Gardner R, Fong CY, Carpenter S, Jardim L, Satishchandra P, Andermann E, Snead OC, 3rd, Lopes-Cendes I, Tsui LC, Delgado-Escueta AV, Rouleau GA, Scherer SW (1998) Mutations in a gene encoding a novel protein tyrosine phosphatase cause progressive myoclonus epilepsy. Nat Genet 20 (2):171–174.

5. Chan EM, Young EJ, Ianzano L, Munteanu I, Zhao X, Christopoulos CC, Avanzini G, Elia M, Ackerley CA, Jovic NJ, Bohlega S, Andermann E, Rouleau GA, Delgado-Escueta AV, Minassian BA, Scherer SW (2003) Mutations in NHLRC1 cause progressive myoclonus epilepsy. Nat Genet 35 (2):125–127.

6. Solaz-Fuster MC, Gimeno-Alcaniz JV, Ros S, Fernandez-Sanchez ME, Garcia-Fojeda B, Criado Garcia O, Vilchez D, Dominguez J, Garcia-Rocha M, Sanchez-Piris M, Aguado C, Knecht E, Serratosa J, Guinovart JJ, Sanz P, Rodriguez de Cordoba S (2008) Regulation of glycogen synthesis by the laforin-malin complex is modulated by the AMP-activated protein kinase pathway. Hum Mol Genet 17 (5):667–678.

7. Kumarasinghe L, Garcia-Gimeno MA, Ramirez J, Mayor U, Zugaza JL, Sanz P (2023) P-Rex1 is a novel substrate of the E3 ubiquitin ligase Malin associated with Lafora disease. Neurobiol Dis 177:105998.

8. Moreno-Estelles M, Campos-Rodriguez A, Rubio-Villena C, Kumarasinghe L, Garcia-Gimeno MA, Sanz P (2023) Deciphering the Polyglucosan Accumulation Present in Lafora Disease Using an Astrocytic Cellular Model. Int J Mol Sci 24 (7):6020.

9. Roma-Mateo C, Aguado C, Garcia-Gimenez JL, Ibanez-Cabellos JS, Seco-Cervera M, Pallardo FV, Knecht E, Sanz P (2015) Increased oxidative stress and impaired antioxidant response in lafora disease. Mol Neurobiol 51 (3):932–946.

10. Ganesh S, Delgado-Escueta AV, Sakamoto T, Avila MR, Machado-Salas J, Hoshii Y, Akagi T, Gomi H, Suzuki T, Amano K, Agarwala KL, Hasegawa Y, Bai DS, Ishihara T, Hashikawa T, Itohara S, Cornford EM, Niki H, Yamakawa K (2002) Targeted disruption of the Epm2a gene causes formation of Lafora inclusion bodies, neurodegeneration, ataxia, myoclonus epilepsy and impaired behavioral response in mice. Hum Mol Genet 11 (11):1251–1262.

11. Criado O, Aguado C, Gayarre J, Duran-Trio L, Garcia-Cabrero AM, Vernia S, San Millan B, Heredia M, Roma-Mateo C, Mouron S, Juana-Lopez L, Dominguez M, Navarro C, Serratosa JM, Sanchez M, Sanz P, Bovolenta P, Knecht E, Rodriguez de Cordoba S (2012) Lafora bodies and neurological defects in malin-deficient mice correlate with impaired autophagy. Hum Mol Genet 21 (7):1521–1533.

12. Garcia-Cabrero AM, Sanchez-Elexpuru G, Serratosa JM, Sanchez MP (2014) Enhanced sensitivity of laforin- and malin-deficient mice to the convulsant agent pentylenetetrazole. Front Neurosci 8:291.

13. Perez-Jimenez E, Viana R, Munoz-Ballester C, Vendrell-Tornero C, Moll-Diaz R, Garcia-Gimeno MA, Sanz P (2021) Endocytosis of the glutamate transporter 1 is regulated by laforin and malin: Implications in Lafora disease. Glia 69 (5):1170–1183.

14. Lahuerta M, Gonzalez D, Aguado C, Fathinajafabadi A, Garcia-Gimenez JL, Moreno-Estelles M, Roma-Mateo C, Knecht E, Pallardo FV, Sanz P (2020) Reactive Glia-Derived Neuroinflammation: a Novel Hallmark in Lafora Progressive Myoclonus Epilepsy That Progresses with Age. Mol Neurobiol 57 (3):1607–1621.

15. Rubio T, Viana R, Moreno-Estelles M, Campos-Rodriguez A, Sanz P (2023) TNF and IL6/Jak2 signaling pathways are the main contributors of the glia-derived neuroinflammation present in Lafora disease, a fatal form of progressive myoclonus epilepsy. Neurobiol Dis 176:105964.

16. Mukhtar I (2020) Inflammatory and immune mechanisms underlying epileptogenesis and epilepsy: From pathogenesis to treatment target. Seizure 82:65–79.

17. Clark IA, Vissel B (2016) Excess cerebral TNF causing glutamate excitotoxicity rationalizes treatment of neurodegenerative diseases and neurogenic pain by anti-TNF agents. J Neuroinflammation 13 (1):236.

18. Bedner P, Steinhauser C (2019) TNFalpha-Driven Astrocyte Purinergic Signaling during Epileptogenesis. Trends Mol Med 25 (2):70–72. doi:S1471-4914(18)30225-30229.

19. Fairless R, Bading H, Diem R (2021) Pathophysiological Ionotropic Glutamate Signalling in Neuroinflammatory Disease as a Therapeutic Target. Front Neurosci 15:741280.

20. Chen Y, Nagib MM, Yasmen N, Sluter MN, Littlejohn TL, Yu Y, Jiang J (2023) Neuroinflammatory mediators in acquired epilepsy: an update. Inflamm Res 72 (4):683–701.

21. Zhang Y, Chen K, Sloan SA, Bennett ML, Scholze AR, O’Keeffe S, Phatnani HP, Guarnieri P, Caneda C, Ruderisch N, Deng S, Liddelow SA, Zhang C, Daneman R, Maniatis T, Barres BA, Wu JQ (2014) An RNA-sequencing transcriptome and splicing database of glia, neurons, and vascular cells of the cerebral cortex. J Neurosci 34 (36):11929–11947.

22. Hardingham GE, Bading H (2010) Synaptic versus extrasynaptic NMDA receptor signalling: implications for neurodegenerative disorders. Nat Rev Neurosci 11 (10):682–696.

23. Naylor DE (2023) In the fast lane: Receptor trafficking during status epilepticus. Epilepsia Open 8 Suppl 1 (Suppl 1):S35–S65.

24. Gallo S, Vitacolonna A, Crepaldi T (2023) NMDA Receptor and Its Emerging Role in Cancer. Int J Mol Sci 24 (3):2540.

25. Rajani V, Sengar AS, Salter MW (2021) Src and Fyn regulation of NMDA receptors in health and disease. Neuropharmacology 193:108615.

26. Dejakaisaya H, Kwan P, Jones NC (2021) Astrocyte and glutamate involvement in the pathogenesis of epilepsy in Alzheimer’s disease. Epilepsia 62 (7):1485–1493.

27. Uddin MS, Lim LW (2022) Glial cells in Alzheimer’s disease: From neuropathological changes to therapeutic implications. Ageing Res Rev 78:101622.

28. Gottlieb M, Matute C (1997) Expression of ionotropic glutamate receptor subunits in glial cells of the hippocampal CA1 area following transient forebrain ischemia. J Cereb Blood Flow Metab 17 (3):290–300.

29. Hogan-Cann AD, Anderson CM (2016) Physiological Roles of Non-Neuronal NMDA Receptors. Trends Pharmacol Sci 37 (9):750–767.

30. Chen TS, Huang TH, Lai MC, Huang CW (2023) The Role of Glutamate Receptors in Epilepsy. Biomedicines 11 (3):783.

31. Huang TH, Lai MC, Chen YS, Huang CW (2023) The Roles of Glutamate Receptors and Their Antagonists in Status Epilepticus, Refractory Status Epilepticus, and Super-Refractory Status Epilepticus. Biomedicines 11 (3):686.

32. Pocock JM, Kettenmann H (2007) Neurotransmitter receptors on microglia. Trends Neurosci 30 (10):527–535.

33. Boileau C, Deforges S, Peret A, Scavarda D, Bartolomei F, Giles A, Partouche N, Gautron J, Viotti J, Janowitz H, Penchet G, Marchal C, Lagarde S, Trebuchon A, Villeneuve N, Rumi J, Marissal T, Khazipov R, Khalilov I, Martineau F, Marechal M, Lepine A, Milh M, Figarella-Branger D, Dougy E, Tong S, Appay R, Baudouin S, Mercer A, Smith JB, Danos O, Porter R, Mulle C, Crepel V (2023) GluK2 Is a Target for Gene Therapy in Drug-Resistant Temporal Lobe Epilepsy. Ann Neurol 94 (4):745–761.

34. Lerma J, Marques JM (2013) Kainate receptors in health and disease. Neuron 80 (2):292–311.

35. Liu J, Feng X, Wang Y, Xia X, Zheng JC (2022) Astrocytes: GABAceptive and GABAergic Cells in the Brain. Front Cell Neurosci 16:892497.

36. Riba M, Auge E, Tena I, Del Valle J, Molina-Porcel L, Ximelis T, Vilaplana J, Pelegri C (2021) Corpora Amylacea in the Human Brain Exhibit Neoepitopes of a Carbohydrate Nature. Front Immunol 12:618193.

37. Singh S, Singh TG, Rehni AK (2020) An Insight into Molecular Mechanisms and Novel Therapeutic Approaches in Epileptogenesis. CNS Neurol Disord Drug Targets 19 (10):750–779.

38. Sherwood MW, Oliet SHR, Panatier A (2021) NMDARs, Coincidence Detectors of Astrocytic and Neuronal Activities. Int J Mol Sci 22 (14):7258.

39. Naylor DE, Liu H, Niquet J, Wasterlain CG (2013) Rapid surface accumulation of NMDA receptors increases glutamatergic excitation during status epilepticus. Neurobiol Dis 54:225–238.

40. Taneja K, Ganesh S (2021) Dendritic spine abnormalities correlate with behavioral and cognitive deficits in mouse models of Lafora disease. J Comp Neurol 529 (6):1099–1120.

41. Sanz P, Garcia-Gimeno MA (2020) Reactive Glia Inflammatory Signaling Pathways and Epilepsy. Int J Mol Sci 21 (11):4096.

42. Munoz-Ballester C, Berthier A, Viana R, Sanz P (2016) Homeostasis of the astrocytic glutamate transporter GLT-1 is altered in mouse models of Lafora disease. Biochim Biophys Acta 1862 (6):1074–1083.

43. Munoz-Ballester C, Santana N, Perez-Jimenez E, Viana R, Artigas F, Sanz P (2019) In vivo glutamate clearance defects in a mouse model of Lafora disease. Exp Neurol 320:112959.

44. Chan KL, Panatpur A, Messahel S, Dahshi H, Johnson T, Henning A, Ren J, Minassian BA (2024) (1)H and (31)P magnetic resonance spectroscopy reveals potential pathogenic and biomarker metabolite alterations in Lafora disease. Brain Commun 6 (2):fcae104.

45. Ikegami A, Haruwaka K, Wake H (2019) Microglia: Lifelong modulator of neural circuits. Neuropathology 39 (3):173–180.

46. Heuser K, Nome CG, Pettersen KH, Abjorsbraten KS, Jensen V, Tang W, Sprengel R, Tauboll E, Nagelhus EA, Enger R (2018) Ca2+ Signals in Astrocytes Facilitate Spread of Epileptiform Activity. Cereb Cortex 28 (11):4036–4048.

47. Dhingra S, Yadav J, Kumar J (2022) Structure, Function, and Regulation of the Kainate Receptor. Subcell Biochem 99:317–350.

48. Negrete-Diaz JV, Falcon-Moya R, Rodriguez-Moreno A (2022) Kainate receptors: from synaptic activity to disease. FEBS J 289 (17):5074–5088.

49. Vargas JR, Takahashi DK, Thomson KE, Wilcox KS (2013) The expression of kainate receptor subunits in hippocampal astrocytes after experimentally induced status epilepticus. J Neuropathol Exp Neurol 72 (10):919–932.

50. Andersen JV, Schousboe A, Wellendorph P (2023) Astrocytes regulate inhibitory neurotransmission through GABA uptake, metabolism, and recycling. Essays Biochem 67 (1):77–91.

51. Molla B, Heredia M, Campos A, Sanz P (2022) Pharmacological Modulation of Glutamatergic and Neuroinflammatory Pathways in a Lafora Disease Mouse Model. Mol Neurobiol 59 (10):6018–6032.

52. Goldsmith D, Minassian BA (2016) Efficacy and tolerability of perampanel in ten patients with Lafora disease. Epilepsy Behav 62:132–135.

53. Zimmern V, Minassian B (2024) Progressive Myoclonus Epilepsy: A Scoping Review of Diagnostic, Phenotypic and Therapeutic Advances. Genes (Basel) 15 (2):171.

