## Supplementary Material for "Glial alterations in the glutamatergic and GABAergic signaling pathways in a mouse model of Lafora disease, a severe form of progressive myoclonus epilepsy"

**Supplementary Table S1:** List of antibodies used in this work, with reference and commercial source.

| **Primary antibody** | **Reference** | **Commercial source** |
|---|---|---|
| Fyn | MA1-19331 | Invitrogen |
| pFyn (Tyr530) | PA5-104756 | Invitrogen |
| GABAA Rα1 | 06-868 | Millipore |
| GABAA Rγ2 | sc-101963 | Santa Cruz Biotech |
| GABA Transporter 1 GAT1 | AGT-001 | Alomone Labs |
| GABA Transporter 3 GAT3 | ab300559 | Abcam |
| GAD65/67 | ab11070 | Abcam |
| GFAP | G3893 | Sigma |
| GluA1 | ab31232 | Abcam |
| GluA2 | ab133477 | Abcam |
| pGluA2 (Ser880) | ab52180 | Abcam |
| GluK2 | ab66440 | Abcam |
| GluN1 | 05-432 | Millipore |
| GluN2A | 07-632 | Millipore |
| GluN2B | MA1-2014 | Invitrogen |
| pGluN2B (Tyr1336) | GTX55068 | GeneTex |
| Iba1 | 234308 | Synaptic Systems |
| mGluR5 | ab76316 | Abcam |
| Lyn | PA5-27361 | Invitrogen |
| pLyn (Tyr397) | bs-3257R | Bioss |
| Src | AHO1152 | Invitrogen |
| pSrc (Tyr416) | 6943 | Cell Signaling |
| Synaptophysin | ab309493 | Abcam |
| vGlut1 | ab272913 | Abcam |
| **Secondary antibody** | **Reference** | **Commercial source** |
| Alexa-Fluor 488 Goat anti Mouse IgG Fc | 115-S45-164 | Jackson Immunoresearch |
| Alexa-Fluor 488 Goat anti Rabbit IgG Fc | 111-S45-046 | Jackson Immunoresearch |
| Alexa-Fluor 568 Donkey anti Mouse IgG (H+L) | A10037 | Thermo Fisher Scientific |
| Alexa-Fluor 568 Donkey anti Rabbit IgG (H+L) | A10042 | Thermo Fisher Scientific |
| Alexa-Fluor 633 Goat anti Guinea Pig IgG (H+L) | A-21105 | Invitrogen |

**Supplementary Table S2: Statistics summary table of the quantification of immunofluorescence and western blot analyses described in this study**. The table details the variable name, the genotype, the number of independent experiments, the intensity of the signal expressed as median ± SD for each variable and the significance (p-value) between control and *Epm2b-/-* samples. An unpaired and non-parametric Mann-Whitney test was performed, according to GraphPad software.

| **Immunofluorescence analyses** | | | |
|---|---|---|---|
| **Variable** | **Genotype** | **Median ± SD** | **p-value** |
| pGluN2B | Control (n: 4) | 13.27 ± 2.58 | 0.0381 |
| | *Epm2b-/-* (n: 6) | 17.41 ± 2.19 | |
| pGluA2 | Control (n: 5) | 25.45 ± 3.02 | 0.111 |
| | *Epm2b-/-* (n: 6) | 29.75 ± 7.26 | |
| GluK2 | Control (n: 5) | 11.41 ± 3.91 | 0.0173 |
| | *Epm2b-/-* (n: 6) | 28.76 ± 16.60 | |
| GAT1 | Control (n: 6) | 16.45 ± 4.50 | 0.0260 |
| | *Epm2b-/-* (n: 6) | 24.78 ± 12.81 | |
| pSrc | Control (n: 5) | 22.38 ± 2.23 | 0.0159 |
| | *Epm2b-/-* (n: 4) | 32.94 ± 4.18 | |
| pLyn | Control (n: 5) | 18.64 ± 3.60 | 0.0317 |
| | *Epm2b-/-* (n: 4) | 30.49 ± 6.40 | |
| **Western blot** | | | |
| **Variable** | **Genotype** | **Mean ± SD** | **p-value** |
| GluN2A | Control (n: 3) | 0.41 ± 0.21 | 0.400 |
| | *Epm2b-/-* (n: 3) | 0.58 ± 0.15 | |
| GluN2B | Control (n: 3) | 0.49 ± 0.11 | 0.400 |
| | *Epm2b-/-* (n: 3) | 0.59 ± 0.02 | |
| pGluN2B | Control (n: 3) | 0.67 ± 0.19 | 0.700 |
| | *Epm2b-/-* (n: 3) | 0.69 ± 0.17 | |
| GluA1 | Control (n: 3) | 1.74 ± 0.61 | 0.100 |
| | *Epm2b-/-* (n: 3) | 2.40 ± 0.22 | |
| GluA2 | Control (n: 3) | 2.23 ± 0.35 | 0.999 |
| | *Epm2b-/-* (n: 3) | 2.39 ± 0.44 | |
| pGluA2 | Control (n: 3) | 1.93 ± 0.77 | 0.200 |
| | *Epm2b-/-* (n: 3) | 3.19 ± 0.91 | |
| mGluR5 | Control (n: 3) | 0.88 ± 0.24 | 0.100 |
| | *Epm2b-/-* (n: 3) | 1.91 ± 0.25 | |

| | | | |
|---|---|---|---|
| GABAAα1 | Control (n: 3) | 0.24 ± 0.11 | 0.999 |
| | *Epm2b-/-* (n: 3) | 0.23 ± 0.21 | |
| GABAAγ2 | Control (n: 3) | 0.64 ± 0.20 | 0.700 |
| | *Epm2b-/-* (n: 3) | 0.55 ± 0.31 | |
| GAD65/67 | Control (n: 3) | 0.66 ± 0.13 | 0.200 |
| | *Epm2b-/-* (n: 3) | 0.94 ± 0.11 | |
| GAT1 | Control (n: 3) | 1.75 ± 0.47 | 0.700 |
| | *Epm2b-/-* (n: 3) | 1.07 ± 0.87 | |
| GAT3 | Control (n: 3) | 3.71 ± 2.29 | 0.999 |
| | *Epm2b-/-* (n: 3) | 3.10 ± 2.89 | |
| pSrc | Control (n: 3) | 0.67 ± 0.07 | 0.100 |
| | *Epm2b-/-* (n: 3) | 1.07 ± 0.06 | |
| Src | Control (n: 3) | 0.16 ± 0.13 | 0.400 |
| | *Epm2b-/-* (n: 3) | 0.19 ± 0.11 | |
| pLyn | Control (n: 3) | 0.61 ± 0.24 | 0.400 |
| | *Epm2b-/-* (n: 3) | 0.82 ± 0.22 | |
| Lyn | Control (n: 3) | 1.00 ± 0.34 | 0.400 |
| | *Epm2b-/-* (n: 3) | 1.12 ± 0.32 | |
| pFyn | Control (n: 3) | 2,82 ± 0.68 | 0.700 |
| | *Epm2b-/-* (n: 3) | 3.53 ± 0.31 | |
| Fyn | Control (n: 3) | 1.13 ± 0.38 | 0.700 |
| | *Epm2b-/-* (n: 3) | 1.47 ± 0.43 | |

**Supplementary Figure Legends:**

**Fig. S1:** Immunofluorescence analyses of different NMDA receptor subunits. A) Diagram of the different areas of the hippocampus analyzed in this work; CA1, Cornu ammonis 1; CA3, Cornu ammonis 3; DG, Dentate gyrus; SR, Stratum radiatum (containing the radiatum, the lacunosum-moleculare, and the molecular layers); from BioRender. B) Representative confocal images of the whole hippocampus derived from control and *Epm2b-/-* mice of 16 months of age, labelled by immunofluorescence with only secondary antibody (No primary ab), anti-GluN1, anti-GluN2A and anti-GluN2B antibodies (in green). At least four independent samples from each genotype (males and females) were analyzed in the same way. The scale corresponds to 200 micrometers. The anti-GluN2B antibody cross-reacted with polyglucosans, even in the presence of 400 mM glucose: see details of the dotted areas containing polyglucosans on the right of the corresponding figures.

**Fig. S2:** Immunofluorescence analyses of the GluA1 subunit of the AMPA receptor. A) Representative confocal images of the whole hippocampus derived from control and *Epm2b-/-* mice of 16 months of age, labelled by immunofluorescence with anti-GluA1 antibodies (in green). At least four independent samples from each genotype (males and females) were analyzed in the same way. The scale corresponds to 200 micrometers. B) Similar samples were co-labelled with anti-GluA1 (in green) and anti-synaptophysin (a pre-synaptic marker) (in red) antibodies. Higher magnification of the Cornu ammonis (CA1), Dentate gyrus (DG), and Stratum radiatum (SR) is shown. The scale corresponds to 20 micrometers.

**Fig. S3:** Immunofluorescence analyses of the GluA2 subunit of the AMPA receptor. A) Representative confocal images of the whole hippocampus derived from control and *Epm2b-/-* mice of 16 months of age, labelled by immunofluorescence with anti-GluA2 antibodies (in green). At least four independent samples from each genotype (males and females) were analyzed in the same way. The scale corresponds to 200 micrometers. B) Similar samples were co-labelled with anti-GluA2 (in green) and anti-synaptophysin (a pre-synaptic marker) (in red) antibodies. Higher magnification of the Cornu ammonis (CA1), Dentate gyrus (DG), and Stratum radiatum (SR) is shown. The scale corresponds to 20 micrometers.

**Fig. S4:** Immunofluorescence analyses of the metabotropic mGluR5 receptor. A) Representative confocal images of the whole hippocampus derived from control and *Epm2b-/-* mice of 16 months of age, labelled by immunofluorescence with anti-mGluR5 antibodies (in green). At least four independent samples from each genotype (males and females) were analyzed in the same way. The scale corresponds to 200 micrometers. B) Similar samples were co-labelled with anti-mGluR5 (in green) and anti-synaptophysin (a pre-synaptic marker) (in red) antibodies. Higher magnification of the Cornu ammonis (CA1), Dentate gyrus (DG), and Stratum radiatum (SR) is shown. The scale corresponds to 20 micrometers.

**Fig. S5:** Immunofluorescence analyses of the GABAA-$\alpha$1 subunit of the GABAA receptor. A) Representative confocal images of the whole hippocampus derived from control and *Epm2b-/-* mice of 16 months of age, labelled by immunofluorescence with anti-GABAA-$\alpha$1 antibodies (in green). At least four independent samples from each genotype (males and females) were analyzed in the same way. The scale corresponds to 200 micrometers. B) Similar samples were co-labelled with anti-GABAA-$\alpha$1 (in green) and anti-synaptophysin (a pre-synaptic marker) (in red) antibodies. Higher magnification of the Cornu ammonis (CA1), Dentate gyrus (DG), and Stratum radiatum (SR) is shown. The scale corresponds to 20 micrometers.

**Fig. S6:** Immunofluorescence analyses of the GABAA-$\gamma$2 subunit of the GABAA receptor, and the GAD65/67 marker, respectively. Representative confocal images of the whole hippocampus derived from control and *Epm2b-/-* mice of 16 months of age, labelled by immunofluorescence with anti-GABAA-$\gamma$2 (panel A) or anti-GAD65/67 (panel B) (in green). At least four independent samples from each genotype (males and females) were analyzed in the same way. The scale corresponds to 200 micrometers.

**Fig. S7:** Immunofluorescence analyses of the GAT3 transporter. A) Representative confocal images of the whole hippocampus derived from control and *Epm2b-/-* mice of 16 months of age, labelled by immunofluorescence with anti-GAT3 antibodies (in green). At least four independent samples from each genotype (males and females) were analyzed in the same way. The scale corresponds to 200 micrometers. B) Similar samples were co-labelled with anti-GAT3 (in green) and anti-synaptophysin (a pre-synaptic marker) (in red) antibodies. Higher magnification of the Cornu ammonis (CA1), Dentate gyrus (DG), and Stratum radiatum (SR) is shown. The scale corresponds to 20 micrometers.

**Fig. S8:** Immunofluorescence analyses of different members of the Src protein kinase family. Representative confocal images of the whole hippocampus derived from control and *Epm2b-/-* mice of 16 months of age, labelled by immunofluorescence with anti-Src, anti-Lyn, anti-Fyn, and anti-pFyn antibodies (in green). At least four independent samples from each genotype (males and females) were analyzed in the same way. The scale corresponds to 200 micrometers.

**Fig. S9:** Western blot analyses of the levels of the proteins assayed in this work. 35 micrograms of proteins of crude extracts from the hippocampus of three controls (C1-C3) and three *Epm2b-/-* (M1-M3) mice of 16 months of age were analyzed by western blot and immunodetection using the antibodies described in Supplementary Table S1. In some antibodies, we did not detect a band at the expected molecular size (*). A) Glutamatergic signaling-related proteins; B) GABAergic signaling-related proteins; C) Src protein kinase family components. Size markers are indicated on the left. The expected molecular size for each protein is indicated in brackets on the left. The panels on the right indicate the quantification of the intensity of the corresponding protein related to the levels of the housekeeping GAPDH protein. Results are expressed as median with a range of three independent samples and represented as arbitrary units (a.u.). Differences between the groups were analyzed by Mann-Whitney non-parametric t-test (see Supplementary Table2 S2). No major changes in the total levels of any of the assayed proteins were observed. In panel A), there is no quantification for the levels of GluN1 or GluK2 is presented because the antibodies did not detect any band of the expected size. In the case of mGluR5, the antibody brochure indicates a predicted band of 132 kDa but an observed band of 280 kDa; the quantification was made on the large protein forms. In panel B), quantification of the GABAA-$\alpha$1 and GAT1 proteins was performed on the main band detected by the corresponding antibodies.

Fig. S1

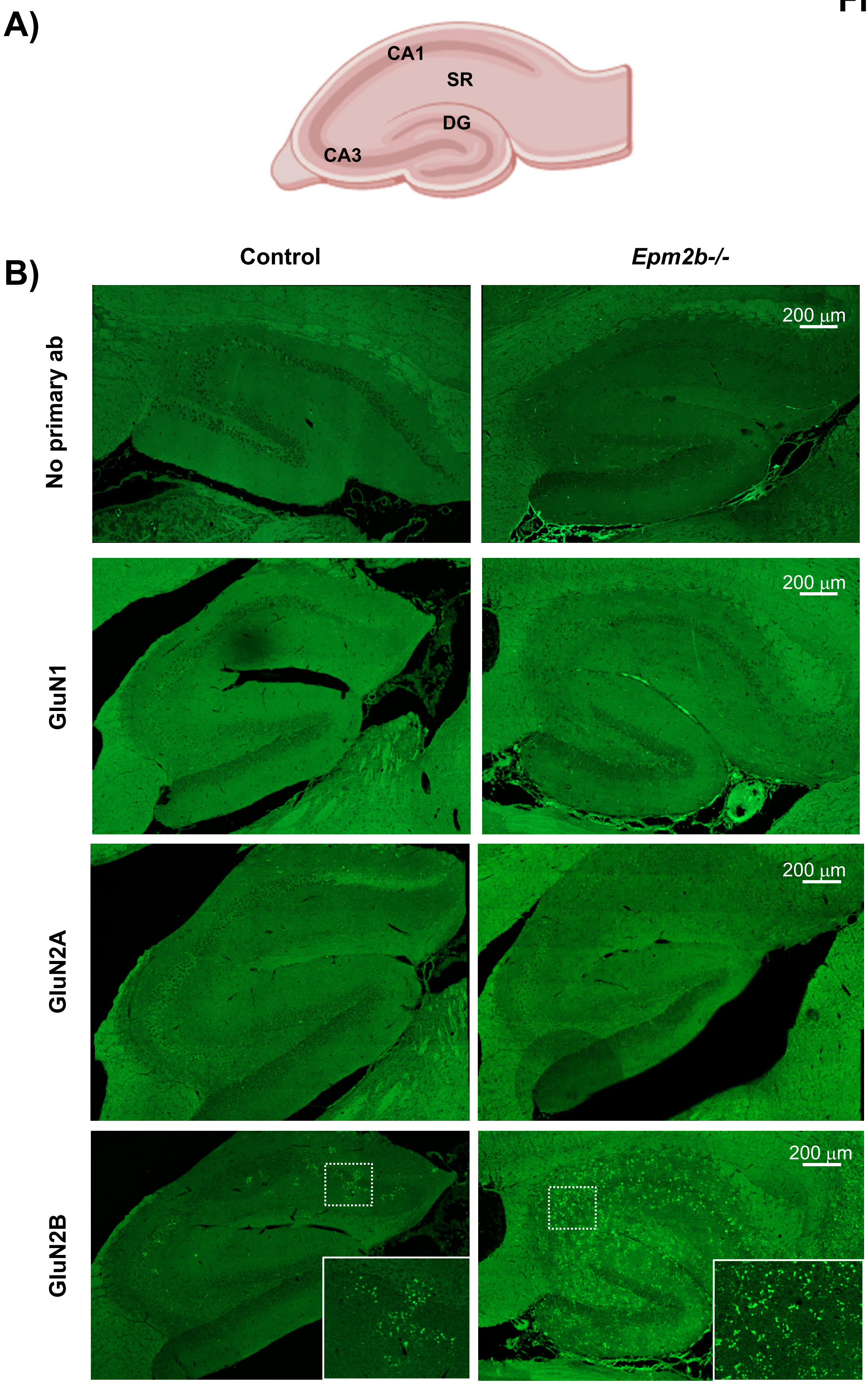
A)
CA1
SR
DG
CA3
B)
Control
Epm2b-/-
No primary ab
GluN1
GluN2A
GluN2B
200 µm
200 µm
200 µm
200 µm

Fig. S2

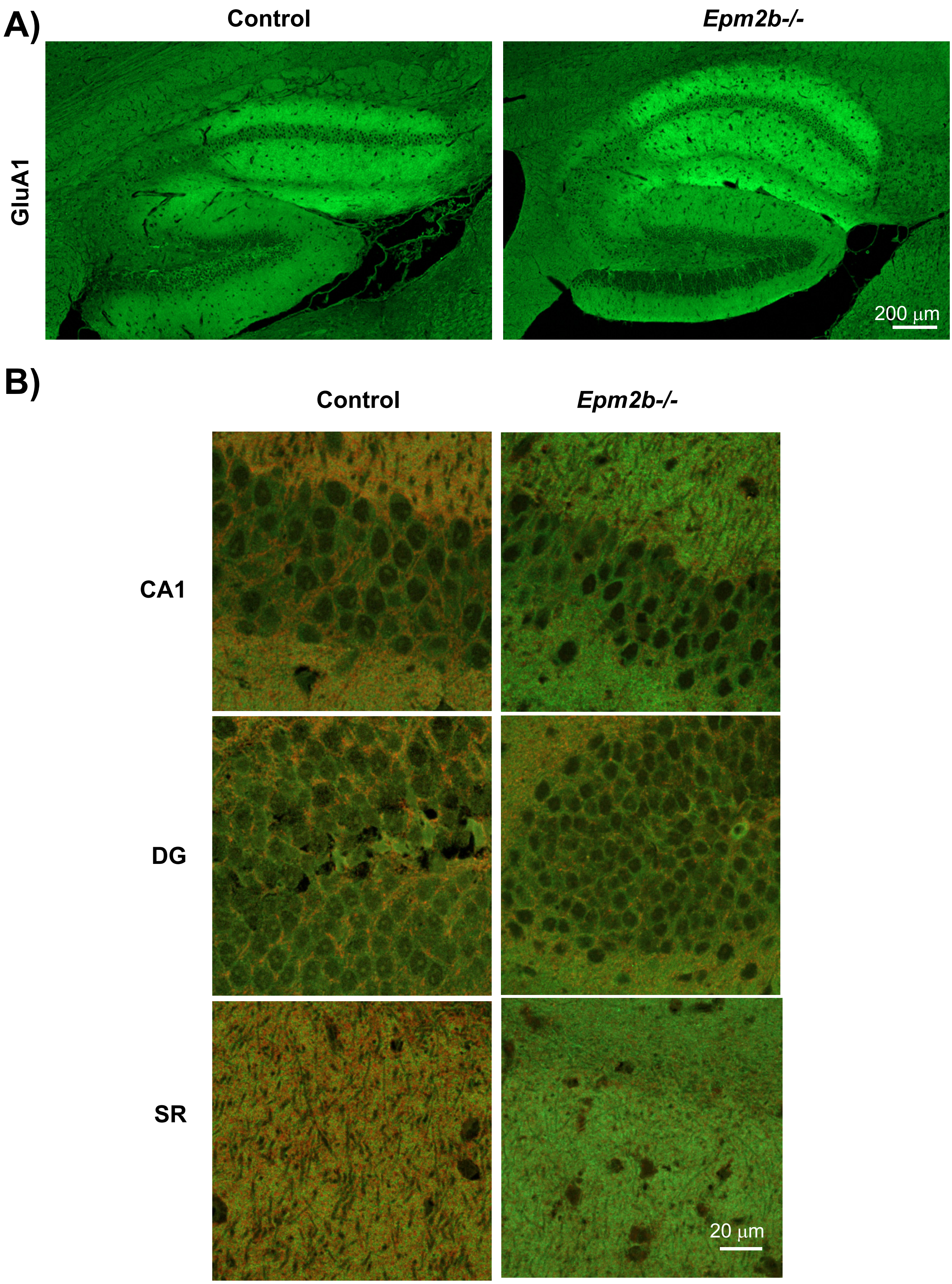

Fig. S3

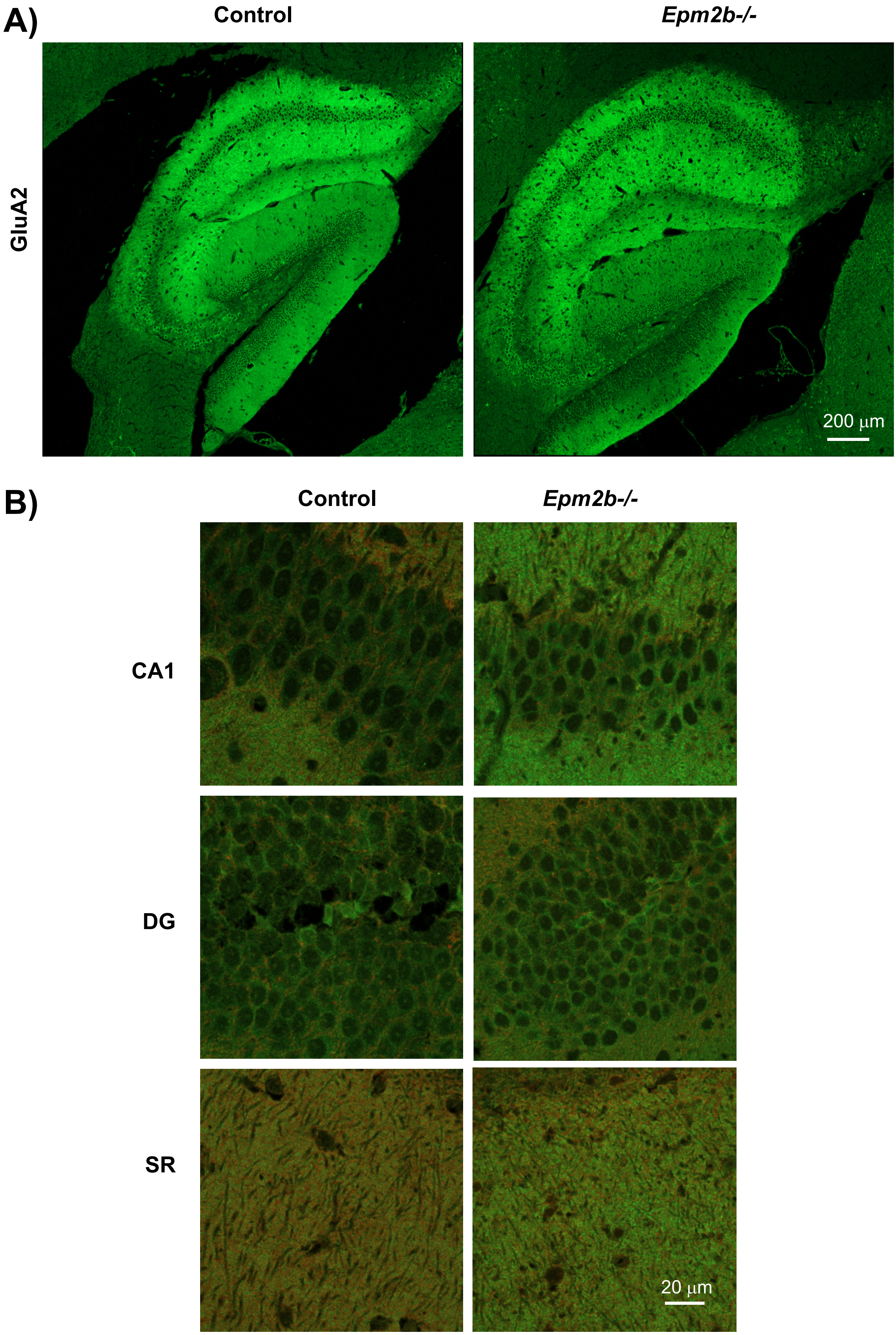

Fig. S4

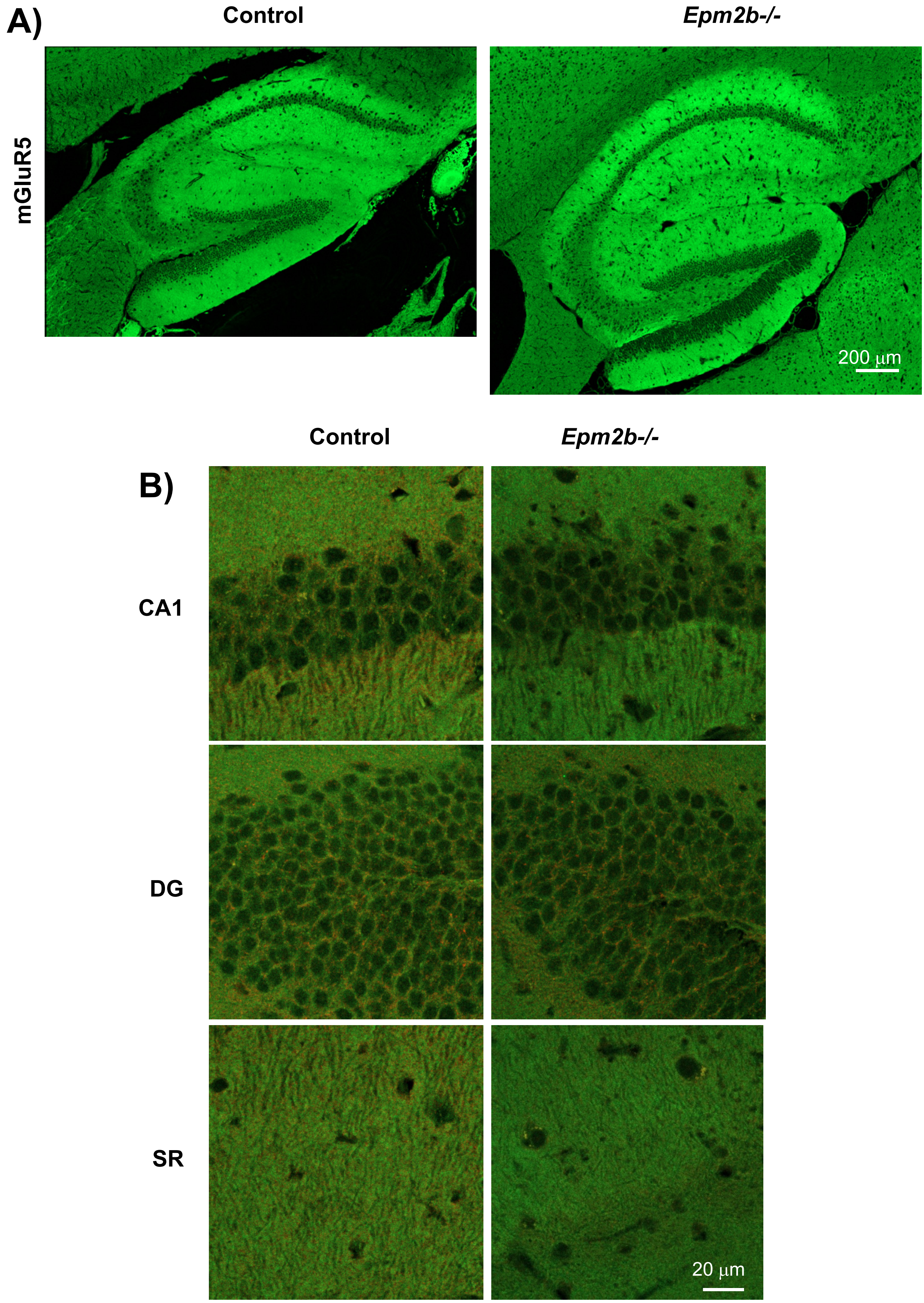

Fig. S5

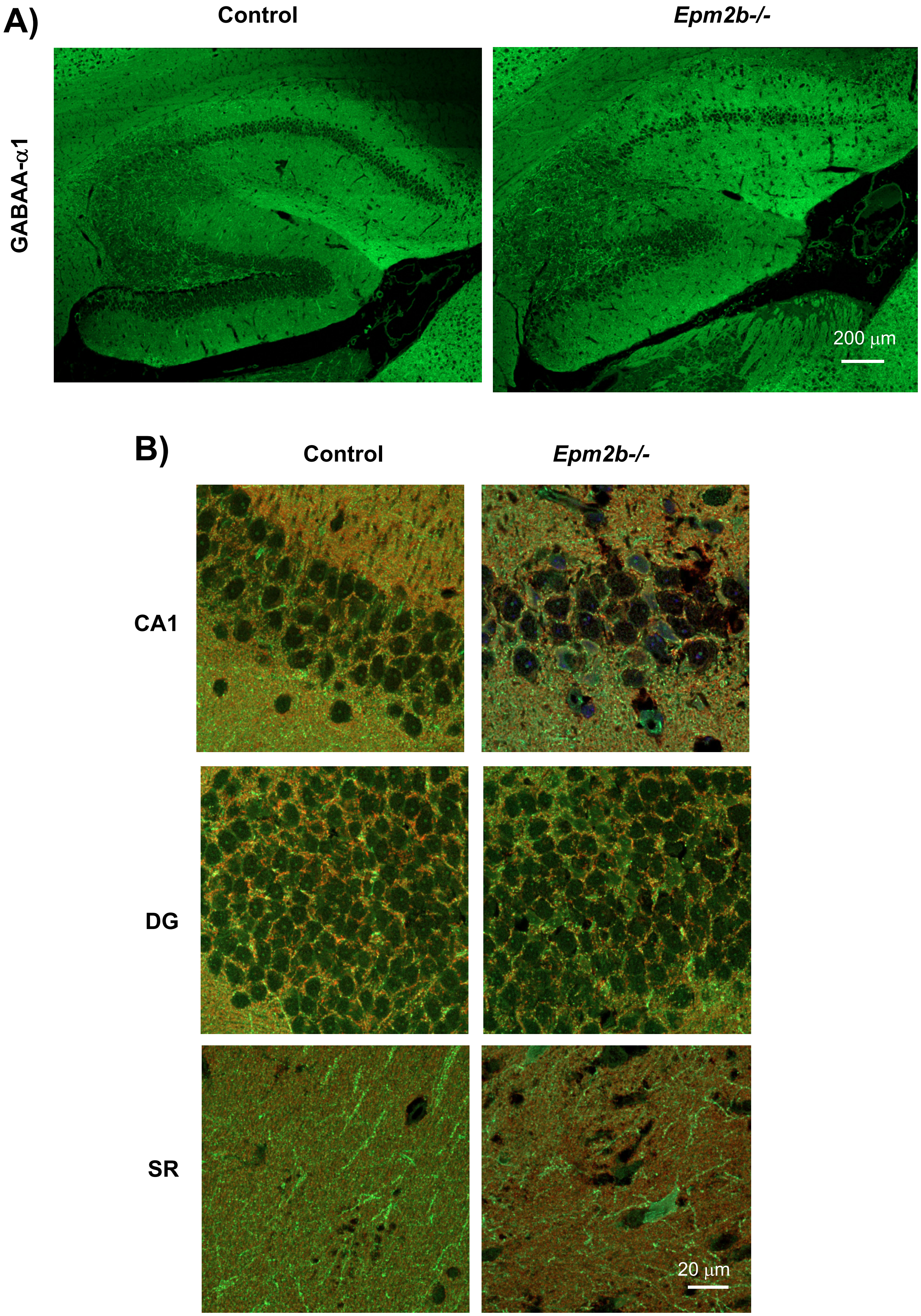

Fig. S6

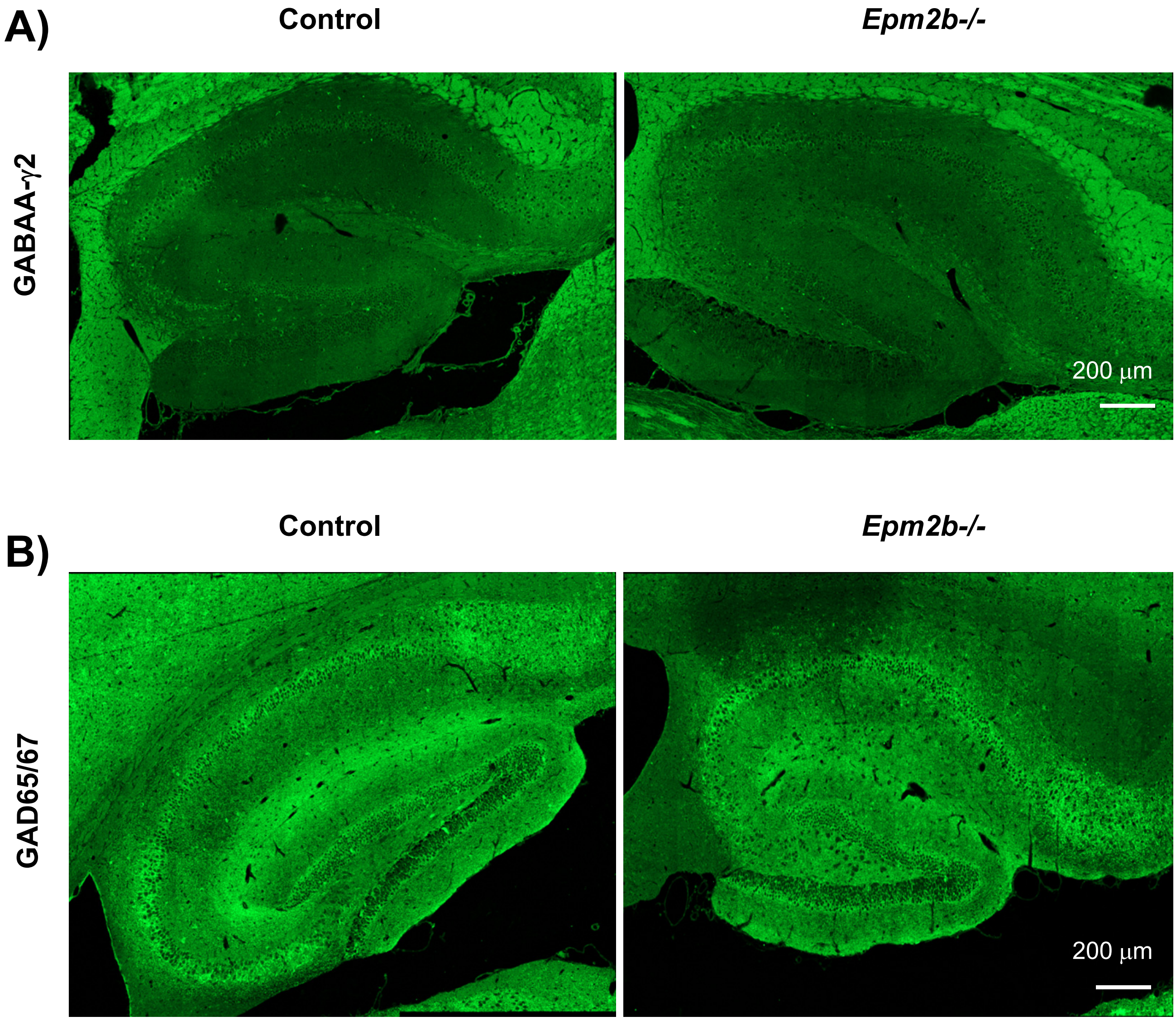
A)
Control
Epm2b-/-
GABAA-γ2
200 µm
B)
Control
Epm2b-/-
GAD65/67
200 µm

Fig. S7

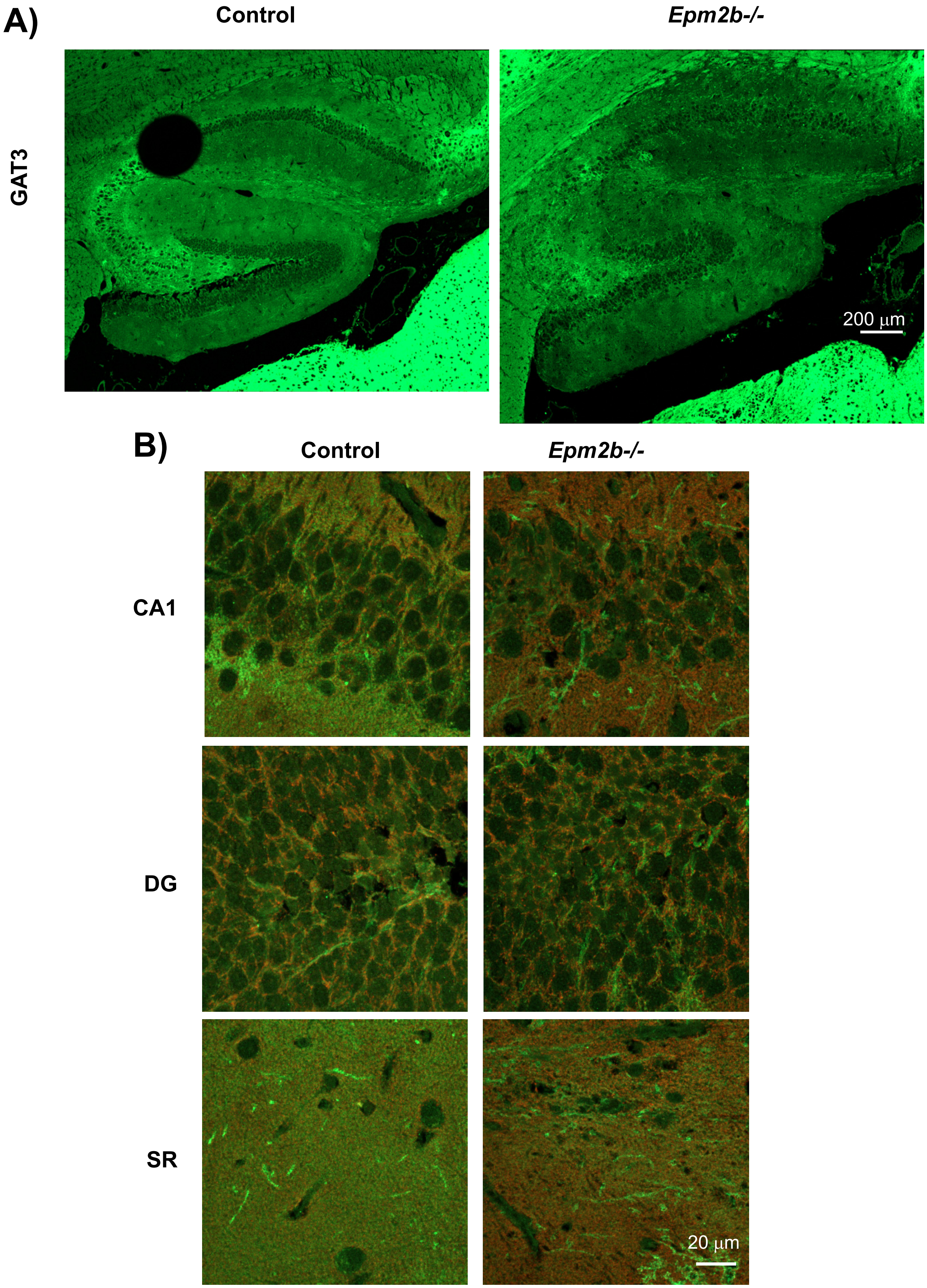

### Fig. S8

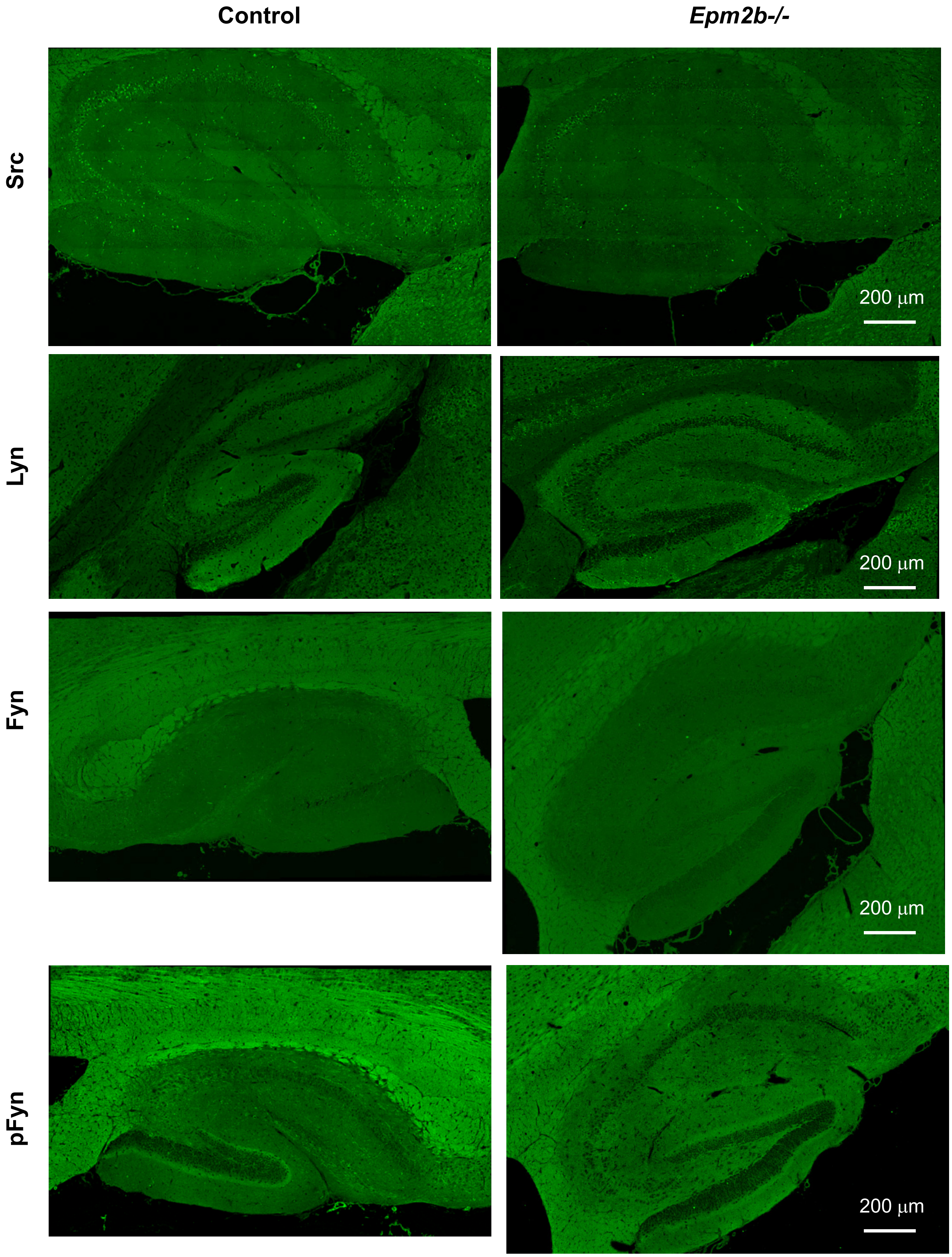

**Fig. S9**

**A)**

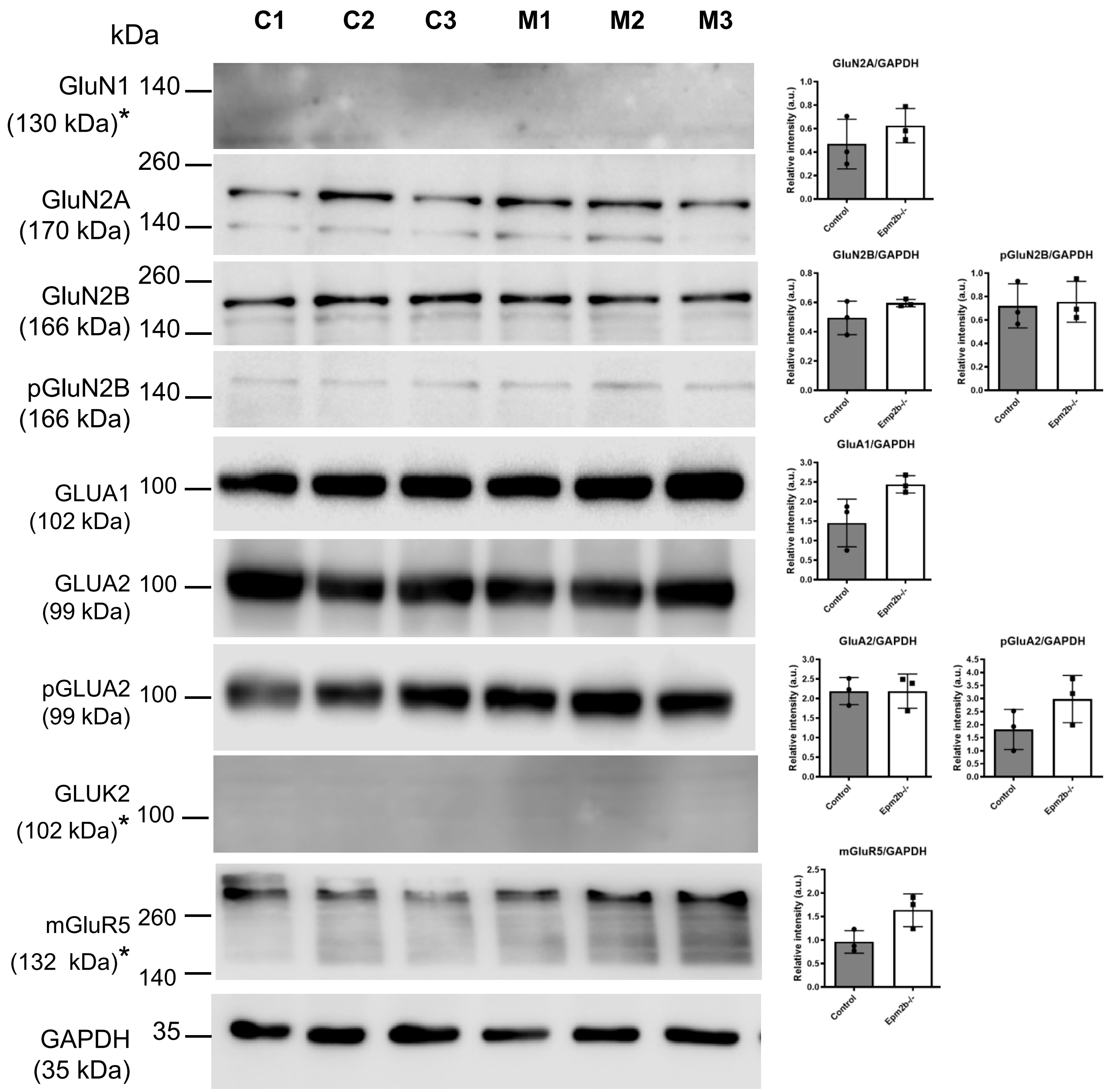

C1
C2
C3
M1
M2
M3
kDa
GluN1 (130 kDa)*
140
GluN2A (170 kDa)
260
140
GluN2B (166 kDa)
260
140
pGluN2B (166 kDa)
140
GLUA1 (102 kDa)
100
GLUA2 (99 kDa)
100
pGLUA2 (99 kDa)
100
GLUK2 (102 kDa)*
100
mGluR5 (132 kDa)*
260
140
GAPDH (35 kDa)
35
GluN2A/GAPDH
GluN2B/GAPDH
pGluN2B/GAPDH
GluA1/GAPDH
GluA2/GAPDH
pGluA2/GAPDH
mGluR5/GAPDH
Relative intensity (a.u.)
Control
Epm2b-/-

### Fig. S9 bis

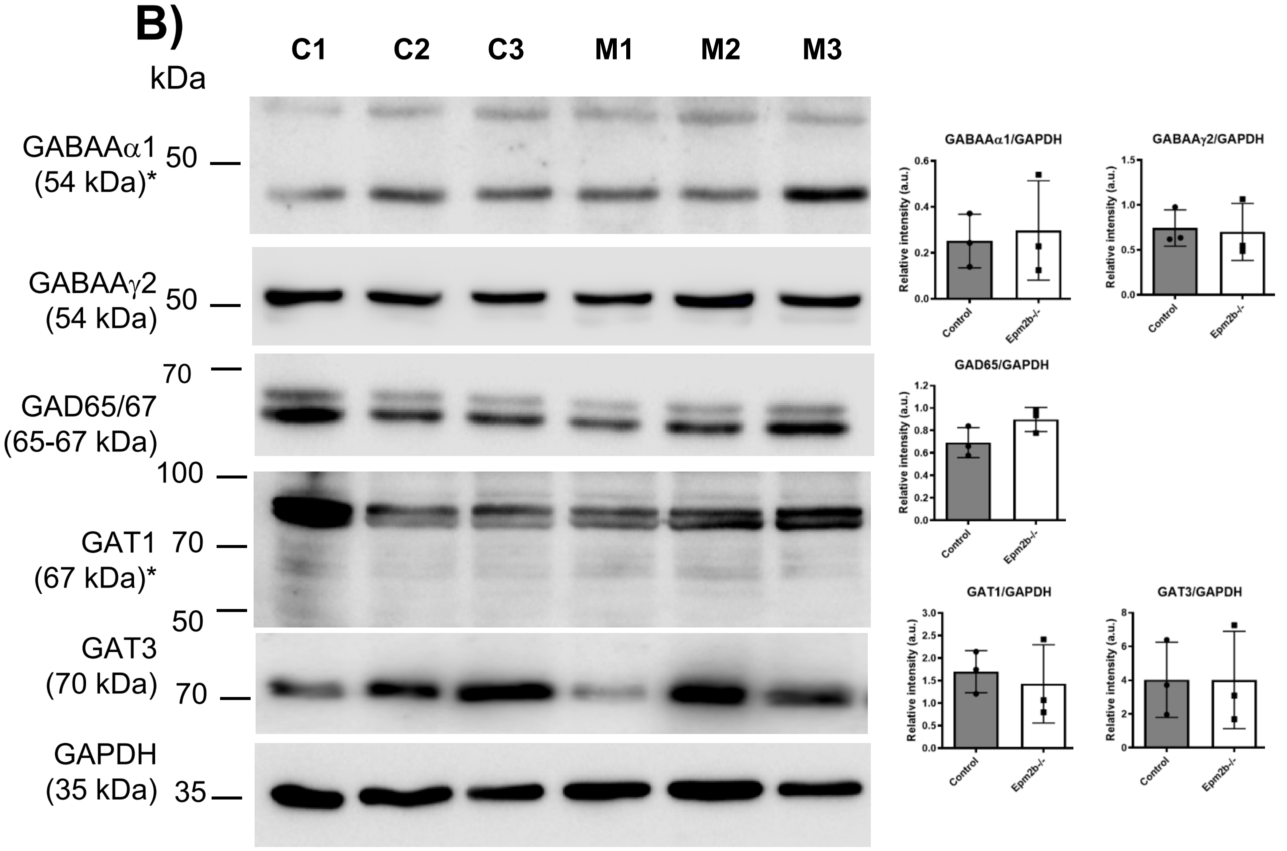
B)
kDa
C1 C2 C3 M1 M2 M3
GABAAα1 (54 kDa)* 50
GABAAγ2 (54 kDa) 50
70
GAD65/67 (65-67 kDa)
100
GAT1 70 (67 kDa)*
50
GAT3 (70 kDa) 70
GAPDH (35 kDa) 35
GABAAα1/GAPDH
GABAAγ2/GAPDH
GAD65/GAPDH
GAT1/GAPDH
GAT3/GAPDH
Relative intensity (a.u.)
Control
Epm2b-/-

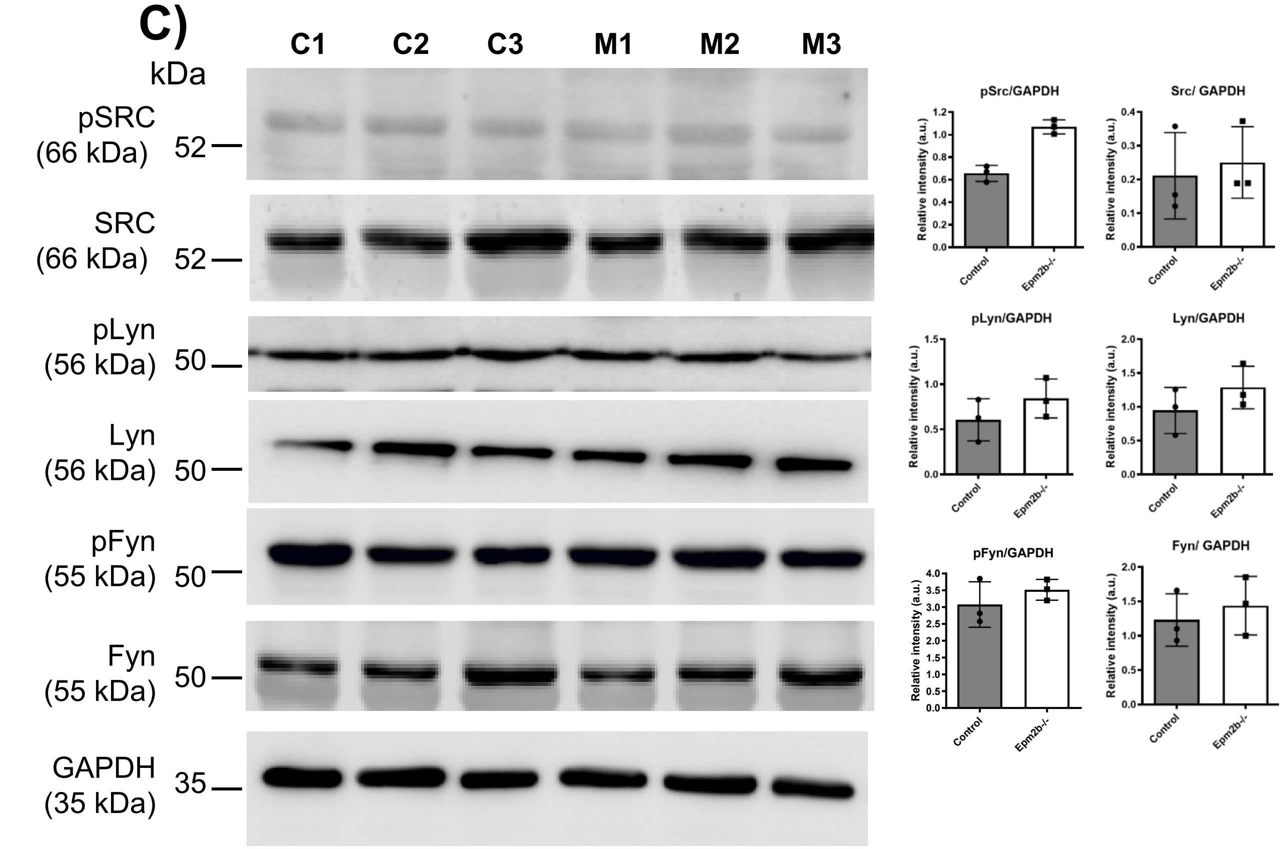
C)
kDa
C1 C2 C3 M1 M2 M3
pSRC (66 kDa) 52
SRC (66 kDa) 52
pLyn (56 kDa) 50
Lyn (56 kDa) 50
pFyn (55 kDa) 50
Fyn (55 kDa) 50
GAPDH (35 kDa) 35
pSrc/GAPDH
Src/ GAPDH
pLyn/GAPDH
Lyn/GAPDH
pFyn/GAPDH
Fyn/ GAPDH
Relative intensity (a.u.)
Control
Epm2b-/-
